# BESTish: A Diffusion-Approximation Framework for Inferring Selection and Mutation in Clonal Hematopoiesis

**DOI:** 10.64898/2026.01.27.702030

**Authors:** Ren-Yi Wang, Khanh N. Dinh, Keito Taketomi, Guodong Pang, Katherine Y. King, Marek Kimmel

**Author notes:** These authors contributed equally to this work (Ren-Yi Wang, Khanh N. Dinh).

## Abstract

Clonal hematopoiesis (CH) arises when hematopoietic stem cells (HSCs) gain a fitness advantage from somatic mutations and expand, resulting in an increase in variant allele frequency (VAF) over time. To analyze CH trajectories, we develop a state-dependent stochastic model of wild-type and mutant HSCs, in which an environmental parameter *α* ∈ [0, 1] regulates death rates and interpolates between homeostatic (Moran-like, *α* = 1) and growth-facilitating (*α* < 1) regimes. Using functional law of large numbers and central limit theorems, we derive explicit mean-field dynamics and a Gaussian–Markov approximation for VAF fluctuations. We show that the mean VAF trajectory has an explicit logistic form determined by selective advantage, while environmental effects affect only the variance and autocovariance structure. Building on these results, we introduce BESTish (Bayesian estimate for selection incorporating scaling-limit to detect mutant heterogeneity), a novel, efficient and accurate Bayesian inference method that can be applied to analyze both cohort-level and longitudinal VAF datasets. BESTish implements the closed-form finite-dimensional distributions that we derive to estimate mutation fitness, mutation rate, and environmental strength for individual CH drivers. When applied to existing CH datasets, BESTish produces consistent mutation fitness inferences across different studies, and estimates CH driver mutation rates in agreement with independent experimental studies. Furthermore, BESTish reveals patient-specific heterogeneity in the selective behavior of recurrent mutations, and identifies variants whose dynamics are compatible with non-homeostatic, growth-facilitating environments. BESTish provides a unified and mechanistic framework for quantifying CH evolution, with potential applications for other biological systems where clonal expansions can be measured.

## 1 Introduction

Hematopoiesis is a highly dynamic and tightly regulated process in which hematopoietic stem cells (HSCs) continuously regenerate the diverse pool of blood and immune cells necessary to sustain human life, dividing to maintain nearly 10^13^ cells distributed throughout the body [Cosgrove et al., 2021]. Despite their central role, the total number of HSCs is relatively small, typically on the order of 10^4^–10^6^ cells in humans [Catlin et al., 2011, Cosgrove et al., 2021, Mitchell et al., 2022], with population-based estimates ranging from approximately 50,000 to 200,000 [Lee-Six et al., 2018]. Moreover, only a fraction of these cells are actively engaged in hematopoiesis at any given time: roughly 30% are thought to be productive [Busch et al., 2015, Cosgrove et al., 2021]. Mitchell et al. [2022] further estimated that 20,000–200,000 cells participate in maintaining blood production, a sum that includes long-, intermediate-, and short-term HSCs as well as multipotent progenitors. With age, HSCs inevitably acquire somatic mutations, some of which provide selective growth advantages. Such mutant clones may expand disproportionately and contribute substantially to hematopoiesis, a phenomenon known as *clonal hematopoiesis* (CH).

Once considered a benign feature of aging, CH is now recognized as a pervasive and clinically relevant phenomenon that bridges somatic evolution, aging, and disease. Large-scale sequencing studies have revealed that CH becomes nearly ubiquitous in older adults and is driven by recurrent mutations in *DNMT3A, TET2, ASXL1*, splicing factors including *SF3B1, SRSF2, U2AF1*, and others [Watson et al., 2020, Fabre et al., 2022, Kar et al., 2022]. These mutations confer fitness advantages to HSCs, leading to exponential clonal expansions whose prevalence and growth rates vary with age.

Studying CH provides a quantitative *in vivo* model for understanding how mutation, selection, and drift shape somatic evolution [Watson et al., 2020]. It also illuminates how aging remodels the hematopoietic landscape and predisposes individuals to disease. Faster-growing clones, particularly those carrying *SRSF2* or *U2AF1* mutations, are associated with increased risk of acute myeloid leukemia and cardiovascular disorders [Fabre et al., 2022]. Environmental and inflammatory factors further modulate CH dynamics, as discussed in [Florez et al., 2022, Hormaechea-Agulla et al., 2021, Bowman et al., 2018, Winter et al., 2024].

The biological system described above can be mathematically represented as a two-compartment model consisting of wild-type (WT) and clonal hematopoiesis (CH) cells. A substantial body of literature addresses mathematical models of multi-compartment cell proliferation, many of which have been applied to the study of cancer initiation and evolution [Durrett, 2015]. In the context of hematopoiesis, spatial constraints within the bone marrow and nonlinear regulatory mechanisms typically limit mathematical analyses to a small number of compartments, requiring the system to be state-dependent. For example, Getto et al. [2013] examined two deterministic models with nonlinear regulation, focusing on global stability. More recently, we extended these models using diffusion approximation techniques to study their stochastic properties [Wang et al., 2025]. Deriving diffusion approximations for the scaled population dynamics greatly facilitates statistical inference, as finite-dimensional distributions can be efficiently computed numerically. An additional advantage of this approach lies in its flexibility: diffusion approximations can be directly mapped to biologically meaningful quantities derived from the underlying population dynamics. This feature is particularly valuable in the study of clonal hematopoiesis, where empirical observations are typically reported as *variant allele frequency* (VAF), approximately half of the ratio between the CH cell count and the total number of hematopoietic stem cells (HSCs).

We develop BESTish (Bayesian estimate for selection incorporating scaling-limit to detect mutant heterogeneity), a quantitative method to infer the mutation and selection rates for individual mutations from their observed VAFs. BESTish is able to infer from both patient-specific longitudinal data, e.g. in [Fabre et al., 2022], and cohort-level observations such as [McKerrell et al., 2015] and [Coombs et al., 2017]. To our knowledge, BESTish represents the first computational effort that utilizes both data types within a consistent mathematical framework to analyze CH mutations, which enhances its applicability for future studies.

BESTish assumes a state-dependent multi-type branching process to model HSC dynamics. The model interpolates between the Moran process (where the population size is conserved; cf. [Wodarz and Komarova, 2014]) and the pure-birth process (where the population size only increases), with the death rate regulated by a parameter *α* ∈ [0, 1] that captures the influence of environmental factors on population growth. We further derive the meanfield approximation of the scaled process as the initial population size becomes large. The mean-field dynamics is deterministic with conserved mass when *α* = 1 and grows exponentially when *α* < 1, which can be used to model dynamics under growth-facilitating conditions (e.g., inflammation). We then recover stochasticity by analyzing the fluctuations around the mean-field dynamics, which can be approximated by a Gauss–Markov process for large initial population size, and we derive explicit expressions for its mean and autocovariance functions, which will serve as a key component in BESTish.

The population-level results culminate in the *variant allele frequency* (VAF) [Dentro et al., 2017], defined as one-half times the proportion of CH cells given that CH mutations are invariably heterozygous. VAF serves as a surrogate for clone size, which is difficult to obtain from biological experiment. We derive explicit expressions for the mean-field approximation of the VAF and characterize the corresponding fluctuation dynamics. Notably, we find that the mean-field trajectory of the VAF follows a logistic curve, independent of *α*, whereas the dependence on *α* arises solely through the fluctuation structure. This result indicates that environmental regulation influences only the second-order characteristics of the VAF dynamics, such as the autocovariance function. Hence, the impact of *α* is difficult to detect when sample sizes are small. In contrast, the fitness advantage of CH mutations can be inferred more readily from the VAF, as it determines the slope and trend of the logistic curve. We also note that Fabre et al. [2022] employed logistic functions to fit VAF trajectories and justified this choice via simulations under the Wright–Fisher model. In the present work, we derive the logistic curve directly from the underlying population dynamics under more general conditions that allow for population growth.

BESTish utilizes these mathematical results to infer key parameters characterizing the VAF dynamics. The algorithm is based on finite-dimensional distributions derived from the diffusion approximation. BESTish is capable of inferring mutation fitness, mutation rate, and strength of environmental factor from both cohort-based and patient-specific longitudinal data.

## 2 Results

### 2.1 HSC dynamics N^(*r*)^ modulated by environmental factor *α*

We assume a branching process for the stem cell dynamics in the bone marrow, beginning with *r* wild-type (WT) cells and no cells carrying clonal hematopoiesis (CH) mutations. The process is characterized with 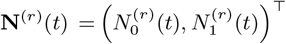, which are respectively the counts of WT and CH cells at time *t*, starting from initial condition **N**^(*r*)^(0) = (*r*, 0). The time until division for cells of type *j* is exponentially distributed with rate *λ*_*j*_ > 0, and WT cells mutate and become CH cells with rate *v*_0_. We assume no further mutation for CH cells, hence *v*_1_ = 0. The impact of the environment factor on the stem cell population is characterized by *α* ∈ [0, 1]. The stochastic dynamics can then be described by the following transitions:

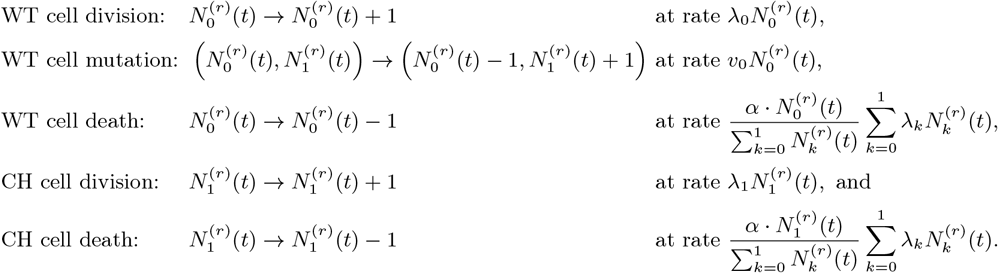

We observe that *α* = 0 correspond to a pure-birth process with mutation, and *α* = 1 is analogous to the Moran process, which will be elucidated in the following section. We define the fitness of type *j* individuals to be *w*_*j*_ := *λ*_*j*_ − *v*_*j*_ and call the CH mutation neutral if *w*_0_ = *w*_1_ and selective if *w*_1_ > *w*_0_. Figure 1A provides a schematic representation of our model, and a rigorous construction of the model with Poisson processes is described in Section A.1 of the Appendix.

**Figure 1.**
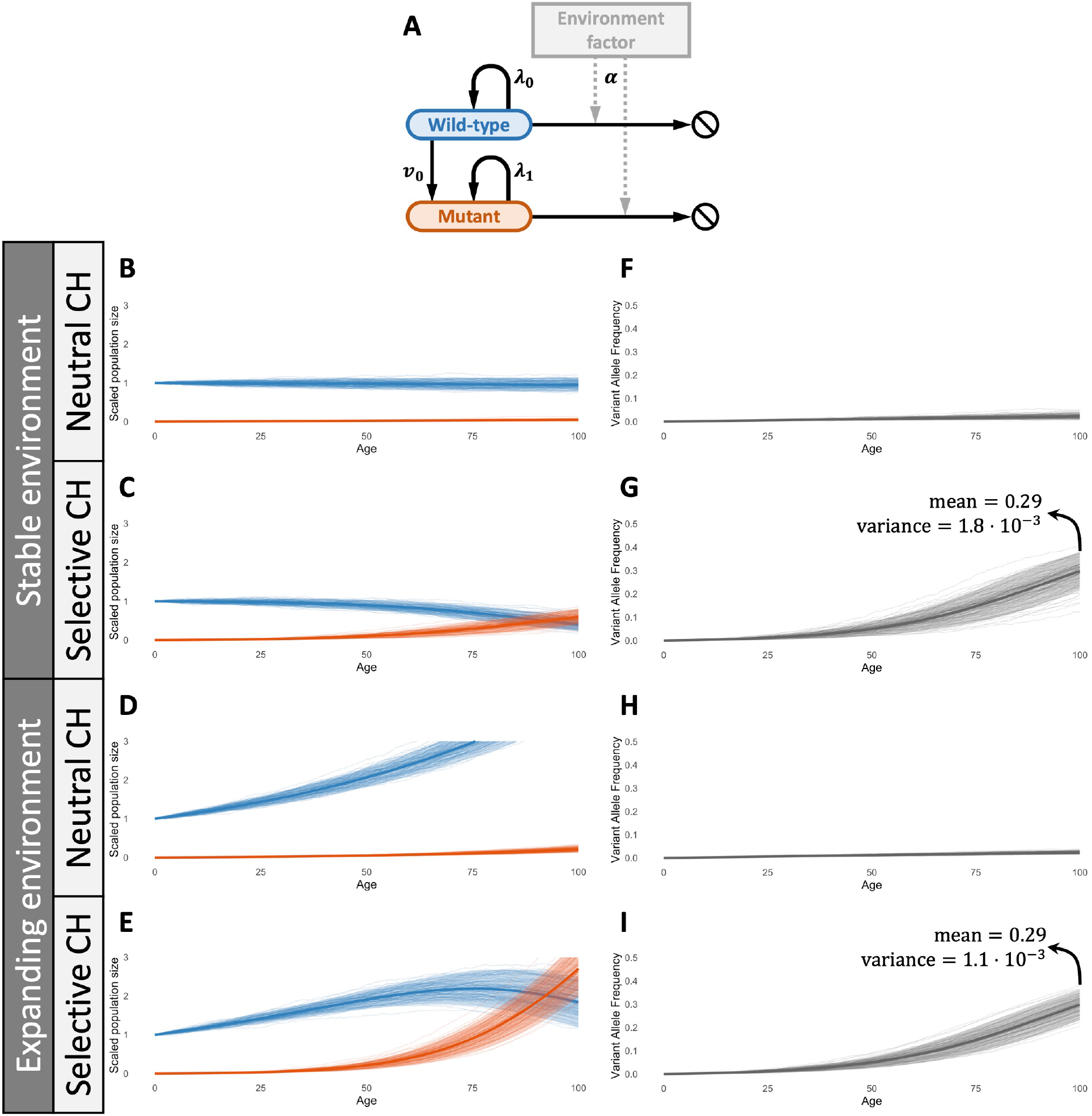
**A**: Schematic for the hematopoietic model with environment factor underlying BESTish. The branching process starts with *r* wild-type cells, which divide at rate *λ*_0_ and mutate at rate *v*_0_. The mutant cells divide at rate *λ*_1_ and cannot mutate (i.e., *v*_1_ = 0). The death rates for both cell types depend on the environment factor, characterized by *α*. **B-I**: Trajectories of scaled population sizes (**B-E**; blue is wild-type, red is mutant) and corresponding variant allele frequency (VAF; **F-I**) in stable (*α* = 1; **B, C, F, G**) and expanding environments (*α* = 0.985; **D, E, H, I**), where the CH mutant is neutral (*w*_0_ = *w*_1_ = 1; **B, F, D, H**) or selective (*w*_0_ = 1, *w*_1_ = 1.05; **C, G, E, I**). Other parameters: *v*_0_ = 5 · 10^−4^, *r* = 20,000. In each plot, expected values (thick lines; Eqs. (1), (2), (7)) and predicted 95%CI regions (shaded areas; Eqs. (6), (10)) are compared against simulations (thin lines).

In Figures 1B-I, we study the simulated dynamics of our HSC model and the resulting VAF trajectories. Figures 1B–E show 100 simulations of the scaled population dynamics 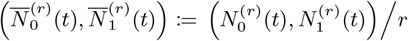, starting from an initial population size of *r* = 20,000 with mutation rate *v*_0_ = 5 *·* 10^−4^. When *α* = 1, we observe conservation of the total scaled population, i.e. 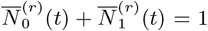, consistent with a homeostatic regime for both neutral mutants (Figure 1B) and selective ones (Figure 1C). In contrast, when *α* < 1, the total population expands exponentially, reflecting growth-facilitating environmental conditions (Figures 1D-E).

Figures 1F–I display the corresponding simulated VAF dynamics, defined as 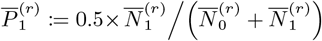. Notably, the mean-field behavior of VAF is invariant with respect to *α*, while the influence of the environmental factor is subtle and manifests only in the fluctuation (e.g., smaller variance in Figure 1I compared to Figure 1G).

In each figure, solid curves and and shaded regions represent the theoretical mean-field approximations and 95% confidence bands respectively, to be derived in the following sections. We observe that the simulations are in agreement with our theoretical results across the parameter space. However, in contrast to simulation-based methods such as approximate Bayesian computation (ABC) [Sisson et al., 2018], BESTish’s parameter inference utilizes the likelihood functions that we derive mathematically in section 2.4. This both ensures the accuracy in BESTish’s parameter estimation and significantly reduces the runtime, as the computational cost for simulations can be extensive, especially for large cell populations.

### 2.2 Mean-field approximation for the population dynamics

We next derive a mean-field approximation for the population dynamics **N**^(*r*)^(*t*). The procedure is analogous to the classical law of large numbers for random variables, treating stochastic processes as random functions. The mean-field approximation captures the central tendency of the dynamics, allowing us to analyze its properties, including growth rate and stability, via differential equations. General discussions on the convergence of stochastic processes can be found at [Billingsley, 1999, Ethier and Kurtz, 2009, Anderson and Kurtz, 2015, Meleard and Bansaye, 2015].

We define the scaled population dynamics to be 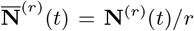, starting from 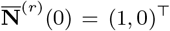. Using the functional law of large numbers (FLLN), we show in Proposition 1 in the Appendix that as *r* → ∞, 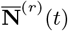 converges almost surely to a deterministic function 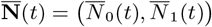, called the *mean-field dynamics* (terminology adapted from [Meleard and Bansaye, 2015]), that follows the initial value problem

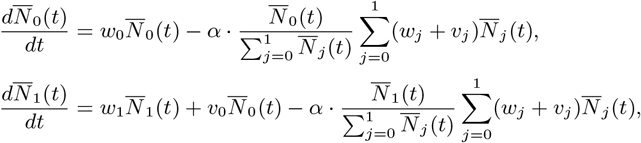

starting from 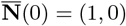, where fitness *w*_*j*_ := *λ*_*j*_ − *v*_*j*_ defines the growth rate of cell type *j*’s population. Details can be found in Proposition 1. Observe that

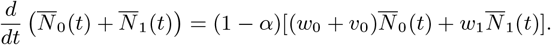

Hence, *α* = 1 leads to the conservation of total population size, where the model is analogous to the Moran process with mutation [Wodarz and Komarova, 2014].

We consider two parameter regimes, one where the CH mutation is neutral (*w*_0_ = *w*_1_) and the mean-field dynamics follows

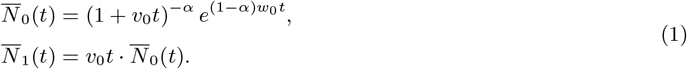

The other case corresponds CH cells are selective (*w*_1_ *> w*_0_) where the dynamics follows

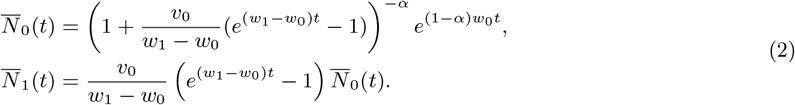

Note that 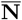 in the neutral regime can be obtained from taking the limit *w*_1_ → *w*_0_ in Equation (2).

### 2.3 Stochastic fluctuation around the mean-field dynamics

In the classical central limit theorem, one takes the scaled difference between the scaled sum of random variables and the mean to recover randomness. Analogously, we center the scaled population dynamics 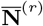 with respect to the mean-field dynamics 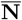 and scale up the difference. The resulting limiting dynamics 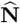 obtained by the functional central limit theorem (FCLT) is called the *fluctuation dynamics*. To outline the procedure, we define the FCLT-scaled dynamics as

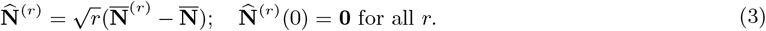

We show in Proposition 2 (Appendix) that as *r* → ∞, 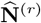 converges to a Gauss-Markov process 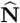 in distribution. Since 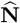 is a Gauss-Markov process, it can be characterized by its mean function **m** and autocovariance function *ρ* (Karatzas and Shreve [2014]). *That is*,

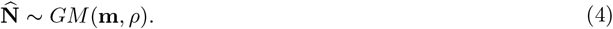

The variance function *V* (*t*) := *ρ*(*t, t*) provides insight into the magnitude of fluctuation of population dynamics 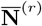 when *r* is large. Specificlly,

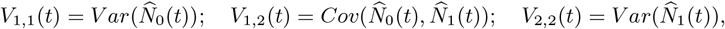

and the autocorrelation function 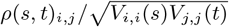 encodes self-dependency and range of dependence of 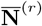. See Wang et al. [2025] for analysis based on convergence of Gaussian processes.

When the initial population size *r* is large, we can decompose the FLLN-scaled dynamics 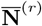 as the mean-field dynamics superposed with scaled fluctuation. Since 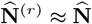 for large *r*, by Eq.(3), we have

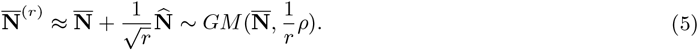

The mean curve and confidence bands for 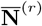 in Figures 1A,C,D,E are computed by Eq.(5) and its direct consequence Eq.(6) below:

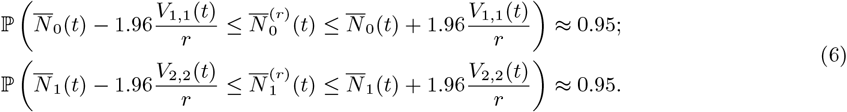

### 2.4 Temporal dynamics of the variant allele frequency of CH cells

The variant allele frequency (VAF) dynamics of a given CH mutation is defined by 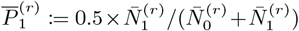 (cf. Dentro et al. [2017]). Using the continuous mapping theorem, we show that 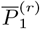 converges almost surely to 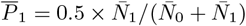 (Theorem 1, Appendix), which can be simplified to

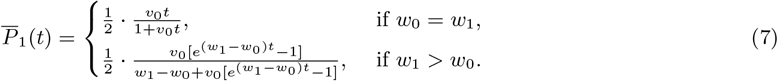

Thus, the mean-field VAF is a logistic curve when the CH mutation confers selective advantage, in accordance to Wright-Fisher simulations in [Fabre et al., 2022]. Our model provides a theoretical derivation for the authors’ observations and generalizes to scenarios with expanding population sizes (when *α* < 1).

The steady state for 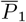 is 1*/*2 for both cases where the CH mutation is neutral (*w*_0_ = *w*_1_) or selective (*w*_1_ *> w*_0_). The asymptotic rate of convergence to the steady state is 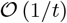 for the neutral scenario. When the CH mutation is selective, 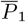 converges to 1*/*2 much faster, at the asymptotic rate 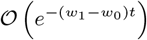 . Hence, it is much harder for neutral mutations to be fixed in a large population. Moreover, 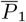 is independent of the environmental parameter *α*, whose effects manifest only at the level of fluctuations. This indicates that VAF is not an ideal surrogate for clone size when the primary objective is to study clonal expansion.

We then employ the delta method to show that as *r* → ∞, 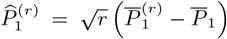 converges in finitedimensional distributions to 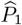 (Theorem 2, Appendix), where

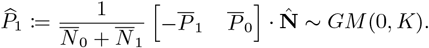

As a result, the autocovariance function *K* for 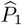 is

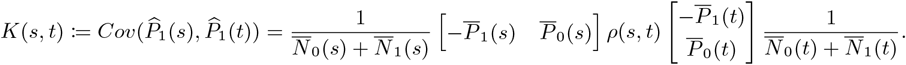

The variance of 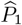 is then

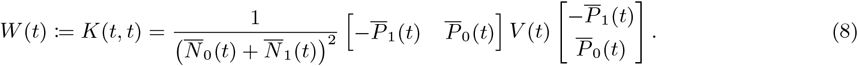

We note that the mean-field VAF 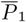 is independent of *α*, while the fluctuation dynamics 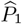 depends on *α*. Similar to Section 2.3, we decompose 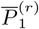 into the deterministic mean-field dynamics and scaled fluctuation for large *r*:

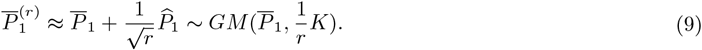

Illustrative longitudinal mean and confidence bands in Figures 1F-I are computed according to Eq. (9) and its direct consequence:

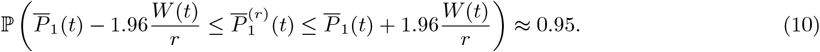

Eq. (9) thus provides us with approximate finite-dimensional distributions for the VAF trajectories, which play a major role in BESTish. To outline the algorithm, we next derive finite-dimensional distributions for both cohort and longitudinal datasets.

### 2.5 A novel algorithm to infer CH mutation parameters from different experimental settings

BESTish estimates the parameters *w*_1_, *v*_0_, and *α* from both cohort-based and patient-specific longitudinal datasets. Since all theoretical results in this work rely on the asymptotic regime, we assume sufficiently large initial population size *r* (typically in the range of 10^4^–10^6^ cells). Finite-dimensional distributions for inference are derived using Eq. (9).

For a cohort dataset 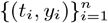 of size *n*, where the CH mutation is observed with VAF = *y*_*i*_ at age *t*_*i*_ in individual *i*, the independence across individuals implies that the joint distribution at times 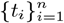 follows

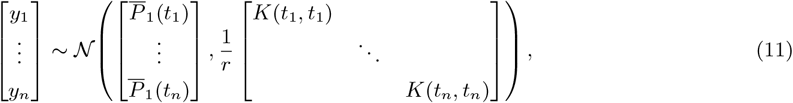

where all off-diagonal entries in the covariance matrix are zero due to the inter-individual independence.

Conversely, for a longitudinal dataset 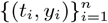 where the CH mutation is observed with VAF = *y*_*i*_ at age *t*_*i*_ in the same individual, the temporal dependence must be retained. In this case, Eq. (9) yields

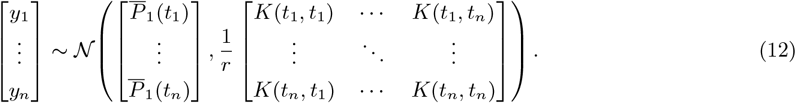

The likelihood functions 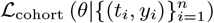 and 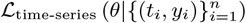 are defined from Eqs.(11) and (12) for a parameter set *θ* using observed VAFs from cohort and time-series datasets, respectively. BESTish infers mutation rate in the logarithmic scale, hence *θ* = (*θ*_1_, *θ*_2_, *θ*_3_, *θ*_4_) = (*w*_0_, *w*_1_, log_10_(*v*_0_), *α*).

We assume that the support of the prior distribution *π*(*θ*) is bounded, i.e. 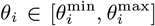. BESTish first divides the range for each parameter *i* into *n*_*i*_ bins, where bin *b* is 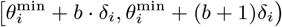. The posterior probability for *θ* in a given bin (*b*_1_, *b*_2_, *b*_3_, *b*_4_) is then

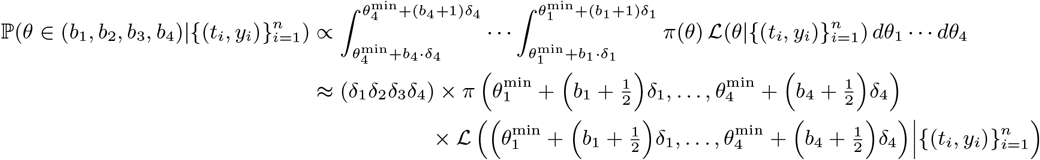

given that the bin sizes *δ*_1_, …, *δ*_4_ are small enough, where ℒ is the appropriate likelihood function for each dataset. BESTish thus computes the posterior probabilities across the *n*_1_ *× · · · × n*_4_ bins, and combines them into the joint posterior distribution across the parameter space.

### 2.6 BESTish reliably infers mutation rate and selective advantage of CH mutations across different data types

We apply BESTish to analyze clonal hematopoiesis data from longitudinal measurements in [Fabre et al., 2022] and cohort-level observations in [McKerrell et al., 2015] and [Coombs et al., 2017]. The model we have employed thus far describes the HSC dynamics within the bone marrow, however the available measurements are derived from peripheral blood samples. Therefore, following Robertson et al. [2022], we assume that the number of differentiated blood cells generated by a given HSC clone is proportional to the size of that clone.

We analyze mutations observed in at least 3 patients in [Fabre et al., 2022] where the variant is measured with VAF *>* 0 at 3 or more distinct time points. Twenty variants in ten genes with longitudinal data are thus studied: *DNMT3A* (*n* = 7 variants), *TET2* (*n* = 3), *SF3B1* (*n* = 2), *SRSF2* (*n* = 2), *CBL* (*n* = 1), *GNB1* (*n* = 1), *IDH2* (*n* = 1), *JAK2* (*n* = 1), *PPM1D* (*n* = 1) and *U2AF1* (*n* = 1). Additionally, cohort data in [McKerrell et al., 2015] and [Coombs et al., 2017] where a variant is observed in ≥ 8 patients are also incorporated in our analysis. This includes *DNMT3A-R882H* (present in *n* = 30 patients in [McKerrell et al., 2015] and *n* = 12 in [Coombs et al., 2017]), *JAK2-V617F* (*n* = 25 in [McKerrell et al., 2015]), *SF3B1-K700E* (*n* = 8 in [McKerrell et al., 2015]) and *SF3B1-K666N* (*n* = 8 in [McKerrell et al., 2015]).

Catlin et al. [2011], Watson et al. [2020], and Mitchell et al. [2022] estimated that HSCs undergo self-renewal approximately once per year, therefore we fix *w*_0_ = 1. For the initial HSC population size, we draw on the estimated number of active HSCs contributing to blood production in [Busch et al., 2015, Cosgrove et al., 2021, Mitchell et al., 2022, Komarova et al., 2024] and set *r* = 20,000. Mitchell et al. [2022] estimated using ABC that the driver mutation rate is 2 *×* 10^−3^ per HSC per year while Watson et al. [2020] estimated the mutation rates to be much lower, on the scale of 10^−6^. For cohort-level inference, we assume uniform prior distributions for *θ* = (*w*_1_, log_10_(*v*_0_), *α*):

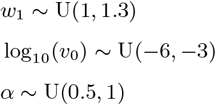

However, given the sparse observations for individual variants in [Fabre et al., 2022], we fix *α* = 1 for longitudinal data to avoid overfitting. We apply BESTish with 500 bins for *w*_1_ and log_10_(*v*_0_), and 100 bins for *α*.

The posterior distributions of (*w*_1_, log_10_(*v*_0_)) for variant *DNMT3A-R882H* from time-series data are presented in Figure 2A. Each patient-specific distribution exhibits a negative correlation between *w*_1_ and log_10_(*v*_0_). This reflects the uncertainty in estimating parameters in our model. The same patient-specific VAF dynamic can be explained by a spectrum of parameters in BESTish: low mutation rate log_10_(*v*_0_) accompanied by high selective advantage *w*_1_, or vice versa.

**Figure 2.**
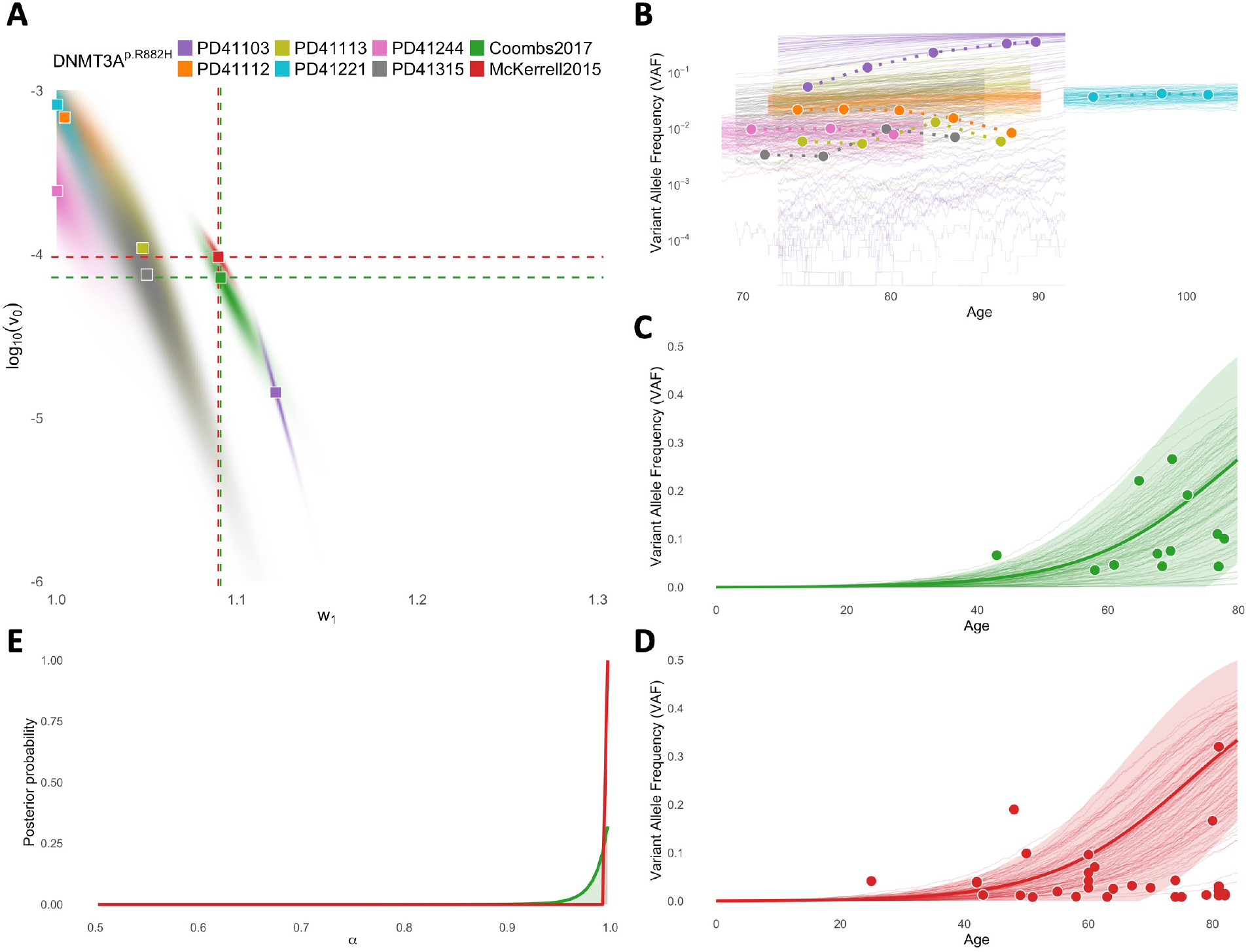
Inference of *DNMT3A-R882H* ‘s mutation rate and selection rate from cohort samples in [Coombs et al., 2017] (“Coombs2017”) and [McKerrell et al., 2015] (“McKerrell2015”), and time-series trajectories in [Fabre et al., 2022] (patient IDs: PD41103, PD411112, PD411113, PD41221, PD41244, PD41315). **A**: Joint posterior distributions for logarithmic mutation rate (log_10_(*v*_0_)) and selection rate (*w*_1_) for each dataset. Squares indicate maximum a posteriori (MAP) estimates from each distribution. Prior distributions: log_10_(*v*_0_) ∼ U(−6, −3), *w*_1_ ∼ U(1, 1.3); for “Coombs2017” and “McKerrell2015”: *α* ∼ U(0.5, 1). Fixed parameters: *w*_0_ = 1, *r* = 2 *×* 10^4^, for time-series data: *α* = 1. **B**: 100 simulated variant allele frequency (VAF) trajectories in logscale assuming MAP estimates (thin lines), against VAFs from corresponding patient-specific longitudinal data (circles, connected by dashed lines). For legibility, simulations are truncated to the age range of the corresponding dataset. **C, D**: Expected values (thick lines) and predicted 95%CI regions (shaded areas) for VAF trajectories assuming MAP estimates, against observed VAFs (circles) and simulations with MAP parameters (thin lines), based on data in [Coombs et al., 2017] (**C**) and [McKerrell et al., 2015] (**D**). **E**: Posterior distributions for environment factor *α* from [Coombs et al., 2017] and [McKerrell et al., 2015]. Colors in **B-E** correspond to datasets in **A**.

Comparing BESTish’s results reveals that *DNMT3A-R882H* behaves more similarly to a neutral mutation (i.e., low *w*_1_) in five patients, and is more characteristic of a driver event (i.e., high *w*_1_) in patient PD41103. Importantly, VAF simulations assuming the maximum a posteriori (MAP) estimates from each distribution closely match the observed dynamics in [Fabre et al., 2022] (Figure 2B), underlying the accuracy of the inferred patient-specific parameters.

The applications of BESTish for *DNMT3A-R882H* using cohort-level measurements separately from [Coombs et al., 2017] and [McKerrell et al., 2015] also result in VAF dynamics that closely match observed data (Figures 2C-D). The expected values and 95%CI regions, derived in previous sections, assuming the MAP estimates for each cohort are also in strong agreement with simulations. Therefore, the implementation of these mathematical results enables BESTish to uncover the parameter distributions more efficiently compared to simulation-based approaches, which would require creating large-scale numerical experiments to approximate the distributions at much higher time costs.

Remarkably, the independent inferences lead to highly consistent results for (*w*_1_, log_10_(*v*_0_)) between the two cohorts (Figures 2A), despite differences in both patient ages ([Coombs et al., 2017]: median = 69 years, range [43, 78]; [McKerrell et al., 2015]: median = 62 years, range [25, 82]) and measured VAFs ([Coombs et al., 2017]: median = 0.072, range [0.035, 0.265]; [McKerrell et al., 2015]: median = 0.027, range [0.008, 0.320]). This confirms that BESTish is robust to data heterogeneity and can reliably infer parameters characterizing individual CH mutations.

We note that the inferred (*w*_1_, log_10_(*v*_0_)) from the cohorts reside within the range of patient-specific inferences from [Fabre et al., 2022] (Figures 2A). This supports our view that cohort-based estimates from BESTish represent population-level averages of the same CH variants’ diverse behaviors in distinct individuals. Finally, the inferred environment factor *α* is approximately 1 in both cohorts (Figures 2E), indicating that the cell dynamics associated with *DNMT3A-R882H* are likely homeostatic.

BESTish’s results from fitting time-series data for 19 other mutations in our study are shown in Supplementary Figures 1-19. Similar to *DNMT3A-R882H*, the simulated VAF trajectories with MAP estimates from distinct inferences closely match the corresponding patient-specific dynamics. This confirms that BESTish can uncover the shared characteristics of a given mutation, despite little overlap in age and VAF ranges from different samples in many instances.

### 3.7 Behavior of CH variants as driver or neutral events is patient-specific

BESTish’s ability to infer patient-specific parameters characterizing a mutation enables a more detailed analysis of CH variants’ diverse behaviors. Figure 3A displays the inferred selection rates for 20 variants across 100 patients included in our study of longitudinal data in [Fabre et al., 2022]. Similar to *DNMT3A-R882H* (Figure 2A), when data in [Coombs et al., 2017] and [McKerrell et al., 2015] is available, the cohort-based inferences consistently fall within the range of estimates derived for individuals (e.g., *JAK2-V617F, SF3B1-K666N* and *SF3B1-K700E*, Figure 3A). Hence, the inter-patient heterogeneity in BESTish’s inferred fitnesses explains the population-level observations for single CH variants.

**Figure 3.**
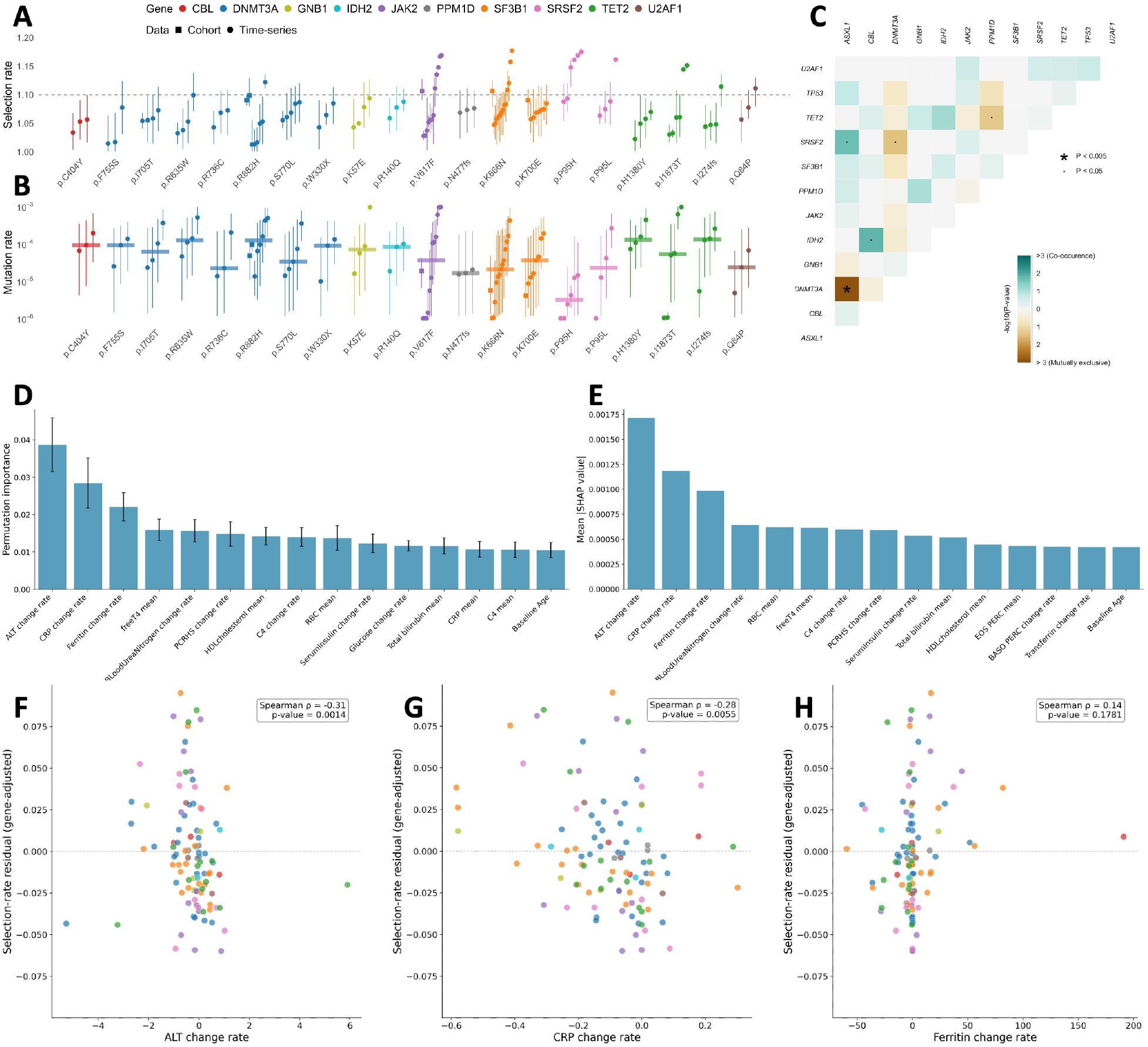
Analysis of mutant-specific intra-patient heterogeneity of fitness advantage inferred from BESTish. **A, B**: Selection rates (**A**) and mutation rates (**B**) inferred from BESTish for 20 frequent CH variants from cohort samples in [McKerrell et al., 2015] and [Coombs et al., 2017] (“Cohort”) and longitudinal trajectories in [Fabre et al., 2022] (“Time-series”). Each point represents the median from the posterior distribution of *w*_1_ (**A**) or log_10_(*v*_0_) (**B**) inferred from a specific cohort or longitudinal patient, and vertical bars display the 95%CI. Horizontal bars in **B** represent the median mutation rates across longitudinal inferences for each CH variant. **C**: Pairwise association matrix between mutated CH genes, using data from [Fabre et al., 2022]. Color indicates the co-occurrence or mutual exclusion of mutations affecting each pair of genes. Gene list consists of *TP53, ASXL1* and those included in **A.**P-values are calculated from Fisher’s exact test, with false discovery rate correction applied to account for multiple testing. **D, E**: Contributions of clinical observations in explaining patient- and variant-specific selection rates in **A**, using feature permutation with random forest (**D**) and SHAP (SHapley Additive exPlanations) (**E**). **F, G, H**: Relationship between BESTish’s inferred variant- and patient-specific selection rates and change rates of ALT (**F**), CRP (**G**) and ferritin (**H**). The color of each mutation (circle) corresponds to genes in **A**. P-value and *ρ* from Spearman’s rank correlation.

By delivering patient-specific parameter estimates, BESTish enables sensitive detection of potential CH driver mutations. The results for *DNMT3A-R882H* (Figure 2) suggest that a threshold of *w*_1_ = 1.1 provides a reasonable criterion for distinguishing neutral from selective CH variants in our model. For *w*_1_ *<* 1.1, the VAF either remains stable or displays negligible growth over time, consistent with neutral evolution (e.g., PD41112, PD411113, PD41221, PD41244, PD41315; Figure 2B). On the other hand, a high selection rate (*w*_1_ *>* 1.1) corresponds to a substantial VAF increase, indicating an ongoing selective sweep (e.g., PD41103; Figure 2B). We identify 8 variants that are highly selective (i.e., *w*_1_ *>* 1.1) in one or more patients: *DNMT3A-R882H, JAK2-V617F, SF3B1-K666N, SRSF2-P95H, SRSF2-P95L, TET2-I1873T, TET2-I274fs* and *U2AF1-Q84P* (Figure 3A). Among these, two variants were overlooked in the original study [Fabre et al., 2022], likely because the authors estimated mutation fitness across multiple patients simultaneously, an approach that is comparable to BESTish’s cohort-based inferences and can only capture the average behaviors of CH variants. The first variant is *I1873T*, determined as likely oncogenic in OncoKB [Chakravarty et al., 2017] and affecting *TET2*, a tumor suppressor and DNA demethylase frequently mutated in hematologic malignancies. The second variant, *P95L* in RNA splicing factor *SRSF2*, is also likely oncogenic [Chakravarty et al., 2017] and has been observed to be a statistically significant hotspot in chronic myeloid leukemia [Zhang et al., 2019].

We then examine the mutation rates inferred from BESTish for the 20 variants in this study (Figure 3B). Despite significant differences in inferred log_10_(*v*_0_) among individuals in the time-series dataset for certain mutations, the median mutation rates across longitudinal patients are always within the 95%CI of the cohort-based inferences when data is available (e.g., *DNMT3A-R882H, JAK2-V617F, SF3B1-K666N* and *SF3B1-K700E*, Figure 3B). Furthermore, the sum of median longitudinal log_10_(*v*_0_) across the 20 variants yields the total mutation rate of 1.2 *×* 10^−3^. This figure is remarkably close to the rate of 2 *×* 10^−3^ driver mutations per HSC per year inferred by Mitchell et al. [2022], who implemented approximate Bayesian computation to estimate mutation rate from single-cell sequencing data for 10 patients between 0 and 81 years of age. The agreement of the inferred driver mutation rates, despite differences in modeling approaches and experimental contexts, further validates the results from BESTish.

We next assess whether the apparent selective behavior of certain CH variants in individual patients may be attributable to other co-occurring driver mutations. To address this, we tabulate the co-occurrence and mutual exclusion in [Fabre et al., 2022] of the 10 genes in this study in addition to *TP53* and *ASXL1*, two genes that have been observed to be frequently mutated in clonal hematopoiesis [Kar et al., 2022](Figure 3C). The only significant pattern is a mutual exclusion between *DNMT3A* and *ASXL1*. There is no significant co-occurrence between other genes, consistent with previous findings by Kar et al. [2022]. This suggests that the CH variants are likely responsible for the selective sweeps observed in patients where BESTish determines their fitness to be high.

We next investigate how clinical covariates contribute to the mutant-specific intra-patient heterogeneity of fitness observed in Figure 3A. We use two approaches, permutation importance and SHAP, to rank the influence of clinical measurements on patient- and mutant-specific selection rates. Because the data are longitudinal with distinct time points for each patient, we summarize each covariate by its mean level and temporal change rate, the latter defined as the slope of the linear regression fitted to measured values against age (details in Methods).

Notably, the three most important covariates identified by both methods are the change rates of alanine aminotransferase (ALT), C-reactive protein (CRP), and ferritin (Figures 3D-E). ALT and CRP change rates correlate with gene-adjusted patient- and mutant-specific selection rates with statistical significance (p-value *<* 0.01, Figures 3F-G), and both proteins are related to liver function and inflammation Wong et al. [2023]. This is consistent with previous findings that CH is associated with elevated pro-inflammatory cytokines, which may act as a driving force for clonal expansion Avagyan and Zon [2023].

We further observe that the mean levels of ALT and CRP are positively correlated with fitness, but these correlations are weaker than those obtained using their change rates. Interestingly, although the three change rates have stronger predictive power for CH mutant fitness, their Spearman rank correlations are negative (Figures 3F-H). This seemingly paradoxical observation requires further investigation, especially given the limited number of individuals with sufficiently dense longitudinal measurements.

### 2.8 The impact of the environment factor

Since *α* < 1 implies net growth of the HSC population, an estimated value of *α* below one suggests that the corresponding mutation may be associated with growth-facilitating conditions. However, when the data are observed only through VAF measurements, *α* is identifiable primarily at the level of fluctuations, which requires substantially more samples for accurate estimation. This limitation arises because VAF is a one-dimensional summary of an underlying two-dimensional process (wild-type and CH cell counts), and thus inevitably entails a loss of information. For instance, illustrative examples in Figures 1C, G, E, I are simulated with identical parameters except for *α*. While the cell population progressions exhibit markedly different behaviors (Figures 1C, E), the corresponding VAF trajectories have identical means throughout time and only differ at the level of fluctuations (Figures 1G, I). This observation further highlights that VAF is not an ideal surrogate for true clone sizes.

In Supplementary Figure 3, the posterior distribution of *α* does not concentrate at 1, suggesting that *SF3B1-K700E* (a splicing-factor mutation) may be associated with growth-facilitating conditions, such as inflammation [Choudhary et al., 2022]. Supplementary Figure 2 shows an even larger deviation of *α* from 1. When combined with the low estimated mutation rate, this pattern suggests that *SF3B1-K666N* may be linked to hematologic malignancy, consistent with clinical observations showing that K666 mutations tend to arrive late and are strongly associated with high-risk MDS and AML [Bick et al., 2020, Chen et al., 2021]. However, these conclusions remain speculative due to limited sample sizes of the cohort-level datasets. We suggest that future studies based on larger population cohorts may utilize BESTish to infer conclusively the role of environmental impacts associated with specific CH variants.

## 3 Discussion

BESTish offers three advantages with respect to parameter inference that set it apart from previous efforts [Zhang and Bozic, 2024, Watson et al., 2020, Fabre et al., 2022]. First, it incorporates our mathematical results for the mean and variance of the VAF dynamics. This enables BESTish to derive the distributions for parameters characterizing each CH variant more accurately and efficiently, compared to simulation-based methods. Second, BESTish can analyze data from either cohort-level studies or time-series experiments. This both enhances its applicability for future studies and allows for consistent analyses of the same CH mutations across different studies and experiments. Finally, BESTish is able to estimate parameters from single patients in longitudinal data, facilitating more in-depth evaluations of covariates influencing CH variants’ heterogeneous behavior as driver or passenger events in different individuals.

Owing to the flexibility of our framework, the methods developed in this paper can be extended to other biological settings involving mutations and interactions with the microenvironment, such as tumorigenesis. A rich literature has proposed stochastic models of tumor evolution with varying levels of mutational complexity [Durrett and Moseley, 2010, Durrett et al., 2011, Antal and Krapivsky, 2011, Nicholson and Antal, 2019, Johnson et al., 2023, Zhang and Bozic, 2024]. However, most existing models do not explicitly incorporate environmental regulation, and their exact branching-process formulations typically render the derivation of finite-dimensional distributions for temporal dynamics analytically intractable. For example, Zhang et al. [2024] emphasize that in multistage tumorigenesis the microenvironment evolves from being tumor-suppressive to malignancy-supporting. In our framework, such a transition can be naturally captured by allowing the environmental parameter *α* to depend on time or on the system state. Moreover, the associated diffusion approximation provides tractable approximate finite-dimensional distributions for the temporal dynamics, thereby substantially simplifying the statistical inference.

## Methods

In this section, we outline step-by-step derivations of quantities that will be used in BESTish. Details and proofs will be deferred to the Appendix.

### State-dependent branching process model

Let 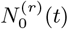 and 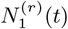 denote the number of WT (type 0) and CH (type 1) cells at time *t* with initial population consisting of *r* WT cells. The rate of division of a type *j* cell is *λ*_*j*_ *>* 0 and the mutation rate is *v*_*j*_ ≥ 0. Since type 1 individuals cannot further mutate, *v*_1_ = 0. All individuals suffer from the same death rate modulated by *α* ∈ [0, 1]:

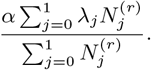

For our model, we define fitness of a type *j* individual by *w*_*j*_ := *λ*_*j*_ − *v*_*j*_ . We are interested in two parameter regimes:

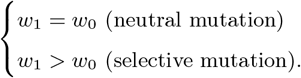

### Functional law of large numbers

Define the density dynamics by

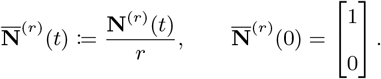

Since *r* represents the initial number of HSCs, it may also be viewed as a proxy for the physical space in which proliferation occurs. Consequently, 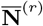 can be interpreted as a density, and increasing *r* corresponds to expanding the effective bone-marrow space.

As *r* → ∞, Proposition 1 shows that 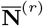 converges in probability to the deterministic limit 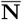, whose components satisfy

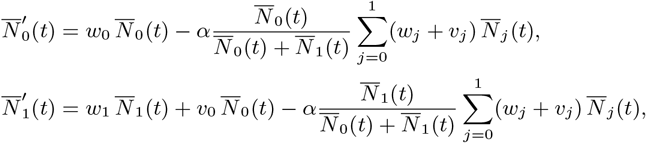

with initial condition 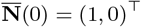.

This is an autonomous system of ODEs, which can be succinctly expressed as 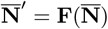.

To solve this system, consider the ratio

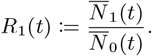

A direct computation shows that the dynamics of *R*_1_ are independent of *α*:

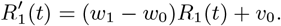

Lemma 1 then gives the explicit solution

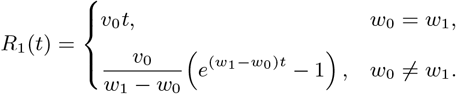

Using 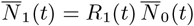, we obtain

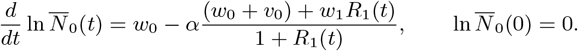

Integrating yields

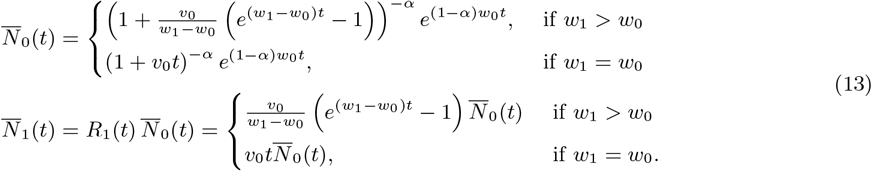

## Functional central limit theorem

Define the fluctuation dynamics by

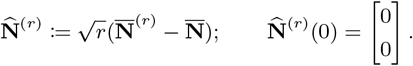

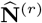 characterizes stochastic fluctuation around the density dynamics. As *r* → ∞, we show in Proposition 2 that 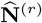 converges in distribution to 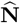, which is characterized by a linear diffusion with time-dependent coefficients

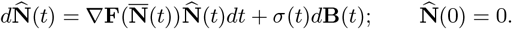

Here, **B** is a 5−dimensional standard Brownian motion and

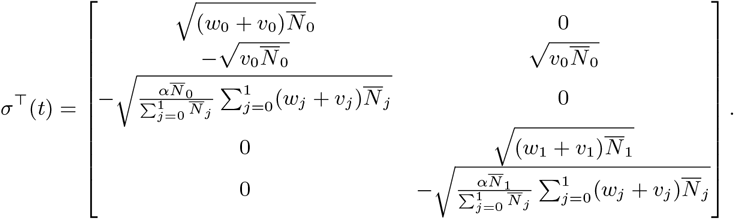

The first row of *σ* corresponds to birth, mutate, and death rates for WT cells and the second row corresponds to birth and death rates for CH cells.

Define a 2 *×* 2 matrix-valued function Φ By

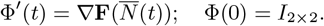

Let Ψ(*t*) := Φ^−1^(*t*). In Proposition 2, we also show that 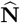 follows a Gauss-Markov process with mean function **m** ≡ 0 and autocovariance function

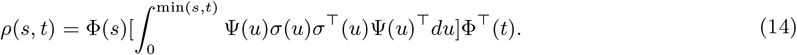

Define the variance function by *V* (*t*) = *ρ*(*t, t*), then

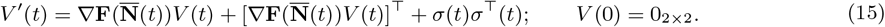

For *s* ≤ *t*, we have

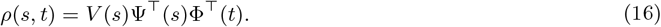

This allows us to compute the joint distribution for patient-specific longitudinal data efficiently in BESTish.

### Variant allele frequency

Define a mapping **g** : 𝒟^2^ → *𝒟*, where 𝒟 is the space of functions that are right-continuous with left limits, such that

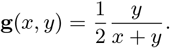

The VAF is then defined by

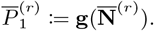

In Theorem 1, we show as *r* → ∞,

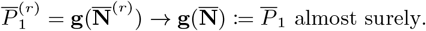

Hence,

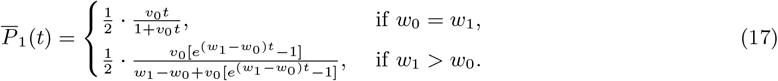

Define 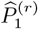 and 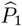 such that for all *t* ≥ 0,

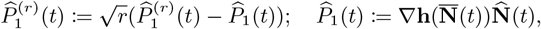

where **h** : ℝ^2^ → ℝ such that

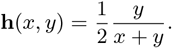

We show using delta method in Theorem 2 that as *r* → ∞, the following convergence in finite-dimensional distribution holds:

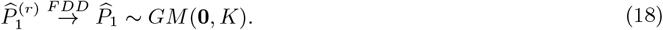

The autocovariance function *K* for 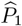 is

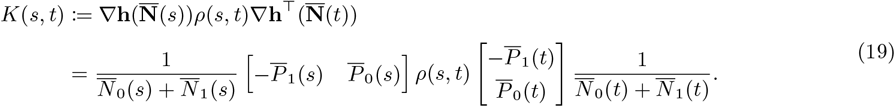

### Finite-dimensional distributions in BESTish

The key ingredient of our algorithm is the Gaussian–Markov approximation

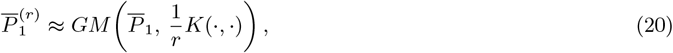

derived from Eq. (18). In other words, 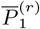 is (approximately) a Gauss-Markov process with mean function 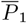 and autocovariance kernel *K/r*. Since the explicit expressions for 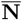 and 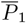 are available (Eqs. (13) and (17)), *K*(*s, t*) in Eq. (19) can be computed by numerical integrating Eq. (14), which can be computed more efficiently by using Eqs. (15) and (16).

For a cohort dataset 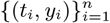 of size *n*, where the CH mutation is observed with VAF = *y*_*i*_ at age *t*_*i*_ in individual *i*, the independence across individuals implies that the joint distribution at times 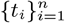 follows

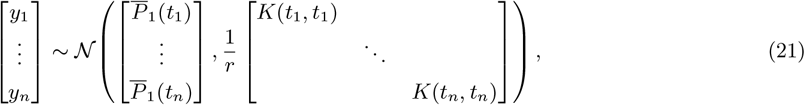

where all off-diagonal entries in the covariance matrix are zero due to the inter-individual independence.

Conversely, for a longitudinal dataset 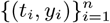 where the CH mutation is observed with VAF = *y*_*i*_ at age *t*_*i*_ in the same individual, the temporal dependence must be retained. In this case, Eq. (20) yields

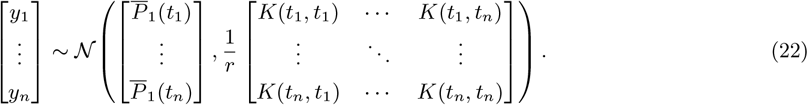

Here, the full covariance matrix captures the temporal correlations in the VAF trajectory.

BESTish infers *θ* = (*θ*_1_, *θ*_2_, *θ*_3_, *θ*_4_) = (*w*_0_, *w*_1_, log_10_(*v*_0_), *α*) using the likelihood functions 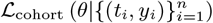 and 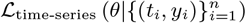, defined based on Eqs. (21) and (22) for population-level and longitudinal data, respectively. Assuming that *θ*_*i*_ is uniformly distributed on 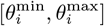, BESTish discretizes the support for each parameter *i* into *n*_*i*_ bins, where bin *b* is 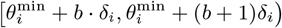. The posterior probability for *θ* in a given bin (*b*_1_, *b*_2_, *b*_3_, *b*_4_) is then

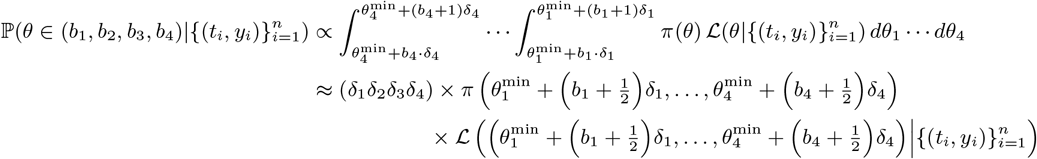

given that the bin sizes *δ*_1_, …, *δ*_4_ are small enough, where ℒ is the appropriate likelihood function for each dataset. BESTish thus computes the posterior probabilities across the *n*_1_ *× · · · × n*_4_ bins, and combines them into the joint posterior distribution across the parameter space.

### Contribution of clinical variates in selection rate heterogeneity

We study the impact of clinical information on the selection rates inferred for the same CH variant across different patients. Longitudinal biological measurements (e.g., in blood) are summarized for each individual with two statistics: mean value and change rate, where the latter is defined as the slope of the linear regression with respect to age.

We separate two sources of heterogeneity in CH fitness: one associated with specific genes and one comprised of other internal and external factors. Comparing the variance of BESTish-inferred CH selection rates and that of the fitness residuals (i.e., selection rates substracted by the gene-specific fitness average) reveals that gene identity explains 24.8% of the selection rate variance in our dataset (100 observations from 89 patients comprising 20 variants across 10 genes).

To analyze the power of clinical information to explain the remaining 75.2% of the fitness variance, we first encode categorical covariates numerically and exclude clinical features with *>* 50% missingness, resulting in a total of 109 clinical covariates. We split the data into a training set (≈ 80% of the observations) and validation set (≈ 20%), such that data from the same patient appears in only one subsample. We train a random forest to predict the fitness residuals from patient-specific clinical covariates using the training set (500 trees; max features = ‘sqrt’; min samples split = 5; min samples leaf = 2), where any missing data is approximated with *k*-nearest-neighbor imputation where *k* = 5. To evaluate the predictive value of the random forest, the same imputation strategy is then performed on the validation set, and model performance is evaluated using 5-fold GroupKFold cross-validation. However, out-of-sample performance for predicting gene-adjusted residuals is low (grouped cross-validation R-squared ≈ −0.09 *±* 0.14), indicating limited out-of-sample predictive performance, likely due to the small sample size. Feature ranking was quantified using permutation importance with 30 repeats, computed as the mean decrease in R-squared after randomly permuting a single feature’s values. However, due to the low predictive power, the feature rankings should be seen as hypothesis-generating rather than predictive. We further complement the random forest approach with SHAP (SHapley Additive exPlanations) values computed via TreeExplainer [Lundberg et al., 2020].

Despite the small sample size, both permutation importance (Figure 3D) and SHAP (Figure 3E) agree on the top-ranking features: change rates of ALT, CRP and ferritin (permutation importance mean *±* standard deviation: 0.0386 *±* 0.0072, 0.0284 *±* 0.0067, and 0.0221 *±* 0.0038, respectively). Spearman rank correlation further confirms that CH selection rates are negatively associated with the change rates of ALT and CRP with statistical significance (*ρ* = −0.31, p-value = 0.0014 and *ρ* = −0.28, p-value = 0.0055, respectively) (Figure 3F-G). In contrast, ferritin change rate is positively associated with increased selection rate but without significance (*ρ* = 0.14, p-value = 0.1781) (Figure 3H).

## Code availability

BESTish is available at https://github.com/dinhngockhanh/BESTish.

## Acknowledgments

RYW, KND and KT acknowledge the support from the Herbert and Florence Irving Institute for Cancer Dynamics at Columbia University. KT is also supported by Krishnan-Ang Philanthropy. RYW, KYK and MK are supported by NIH grant P01CA265748. KYK is also supported by NIH grant R35 HL155672. GP is supported by NSF grants DMS 2216765 and CMMI 2452829.

---

**Supplementary Figure 1:**
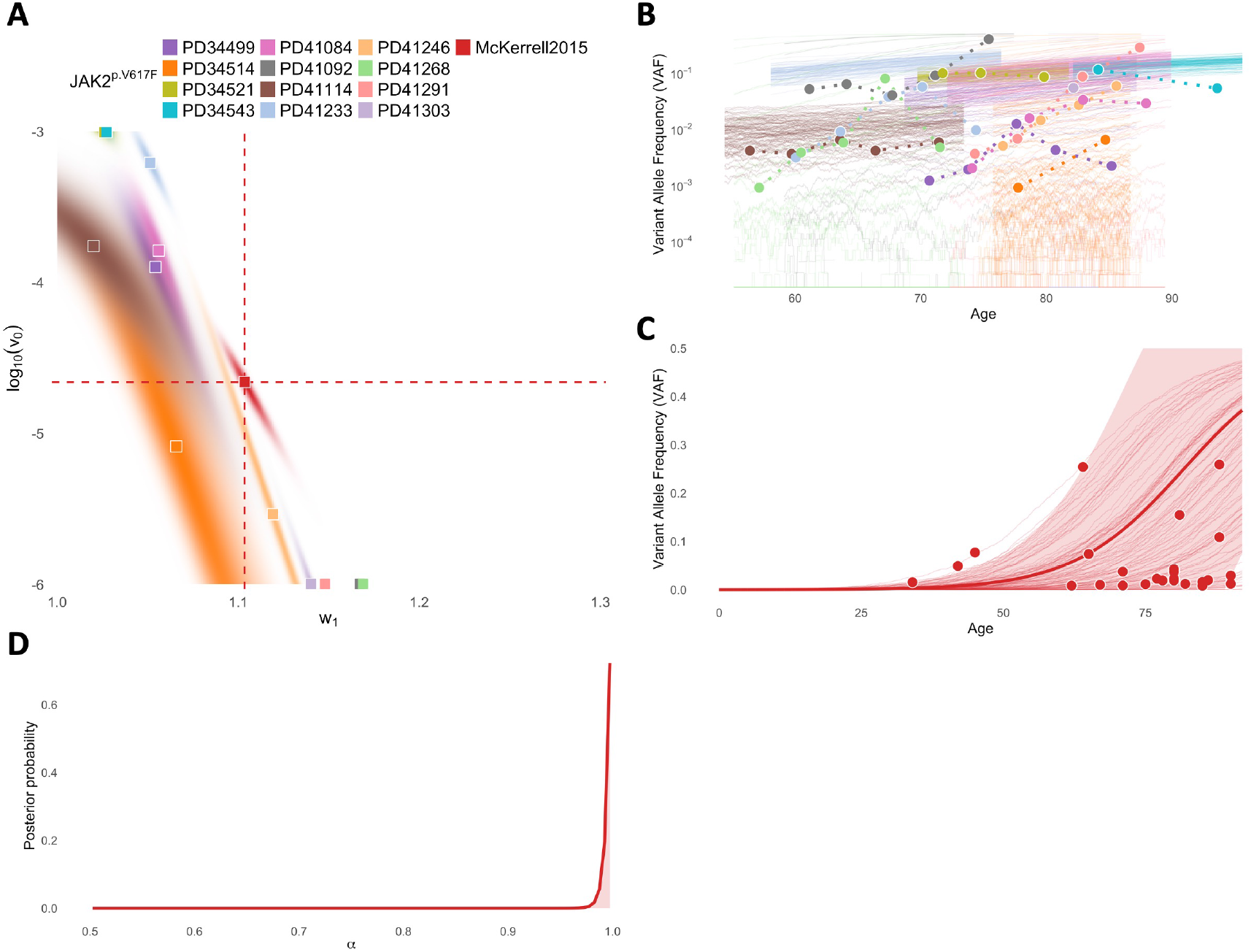
Inference of *JAK2-V617F* ‘s mutation rate and selection rate from cohort samples in [McKerrell et al., 2015] (“McKerrell2015”) and time-series data in [Fabre et al., 2022]. **A**: Joint posterior distributions for log_10_(*v*_0_) and *w*_1_. Squares = MAP estimates. **B**: 100 simulated VAF trajectories in logscale assuming MAP estimates (thin lines), against observed VAFs (circles, connected by dashed lines) for samples in [Fabre et al., 2022]. **C**: Expected values (thick lines) and predicted 95%CI regions (shaded areas) for VAF trajectories assuming MAP estimates, against observed VAFs (circles) and simulations with MAP parameters (thin lines) for samples in [McKerrell et al., 2015]. Colors in **B-D** correspond to datasets in **A**.

**Supplementary Figure 2:**
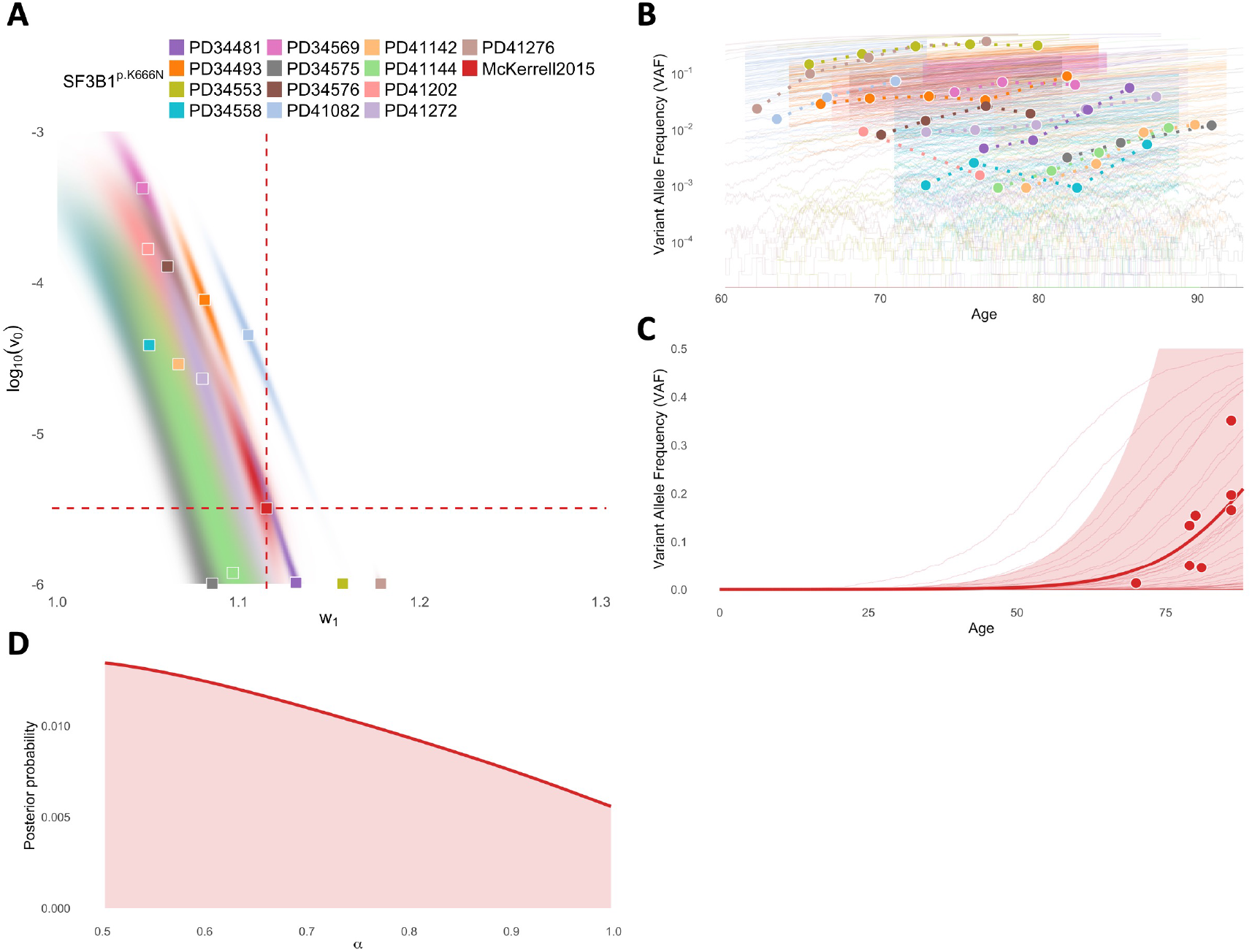
Inference of *SF3B1-K666N* ‘s mutation rate and selection rate from cohort samples in [McKerrell et al., 2015] (“McKerrell2015”) and time-series data in [Fabre et al., 2022]. **A**: Joint posterior distributions for log_10_(*v*_0_) and *w*_1_. Squares = MAP estimates. **B**: 100 simulated VAF trajectories in logscale assuming MAP estimates (thin lines), against observed VAFs (circles, connected by dashed lines) for samples in [Fabre et al., 2022]. **C**: Expected values (thick lines) and predicted 95%CI regions (shaded areas) for VAF trajectories assuming MAP estimates, against observed VAFs (circles) and simulations with MAP parameters (thin lines) for samples in [McKerrell et al., 2015]. Colors in **B-D** correspond to datasets in **A**.

**Supplementary Figure 3:**
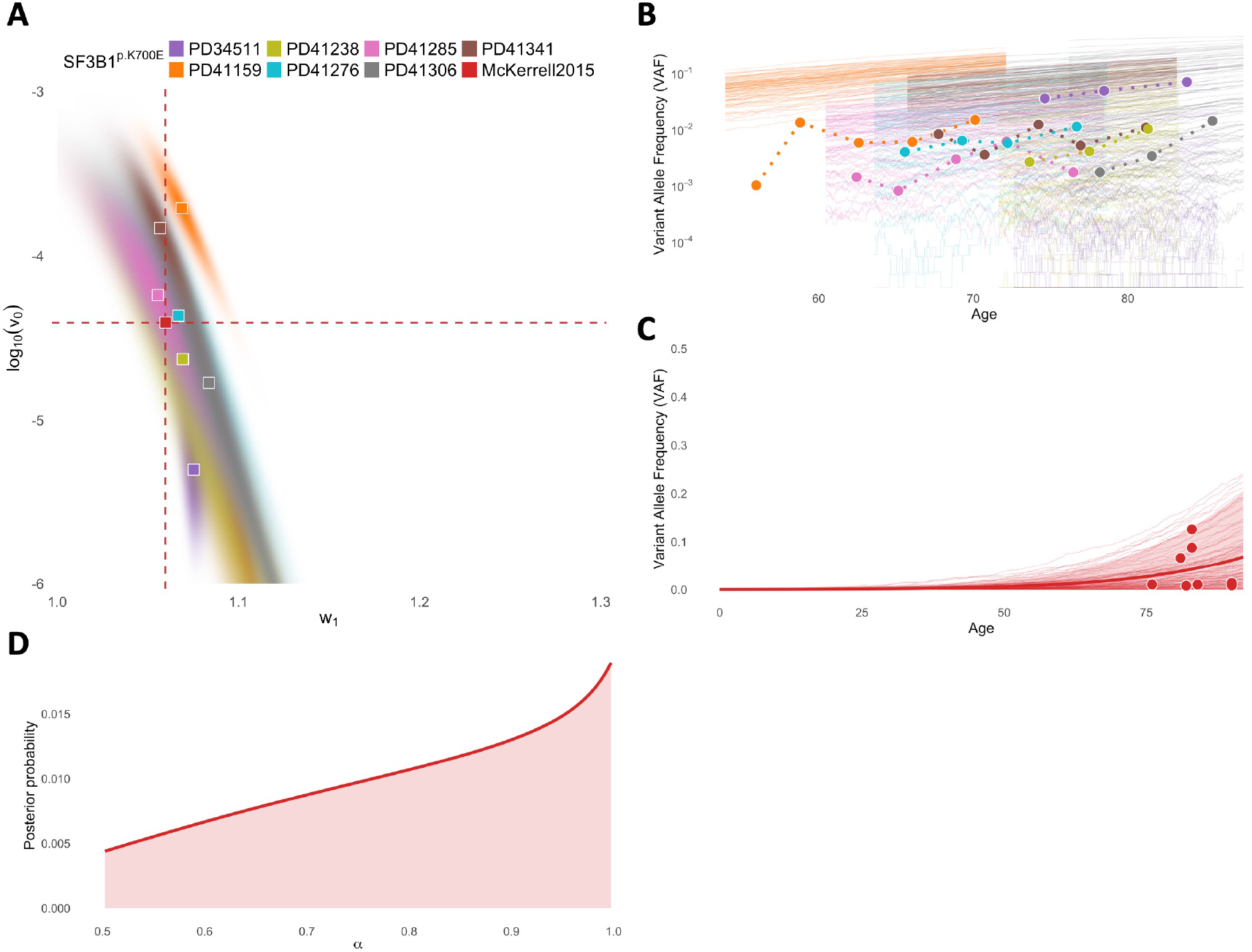
Inference of *SF3B1-K700E* ‘s mutation rate and selection rate from cohort samples in [McKerrell et al., 2015] (“McKerrell2015”) and time-series data in [Fabre et al., 2022]. **A**: Joint posterior distributions for log_10_(*v*_0_) and *w*_1_. Squares = MAP estimates. **B**: 100 simulated VAF trajectories in logscale assuming MAP estimates (thin lines), against observed VAFs (circles, connected by dashed lines) for samples in [Fabre et al., 2022]. **C**: Expected values (thick lines) and predicted 95%CI regions (shaded areas) for VAF trajectories assuming MAP estimates, against observed VAFs (circles) and simulations with MAP parameters (thin lines) for samples in [McKerrell et al., 2015]. Colors in **B-D** correspond to datasets in **A**.

**Supplementary Figure 4:**
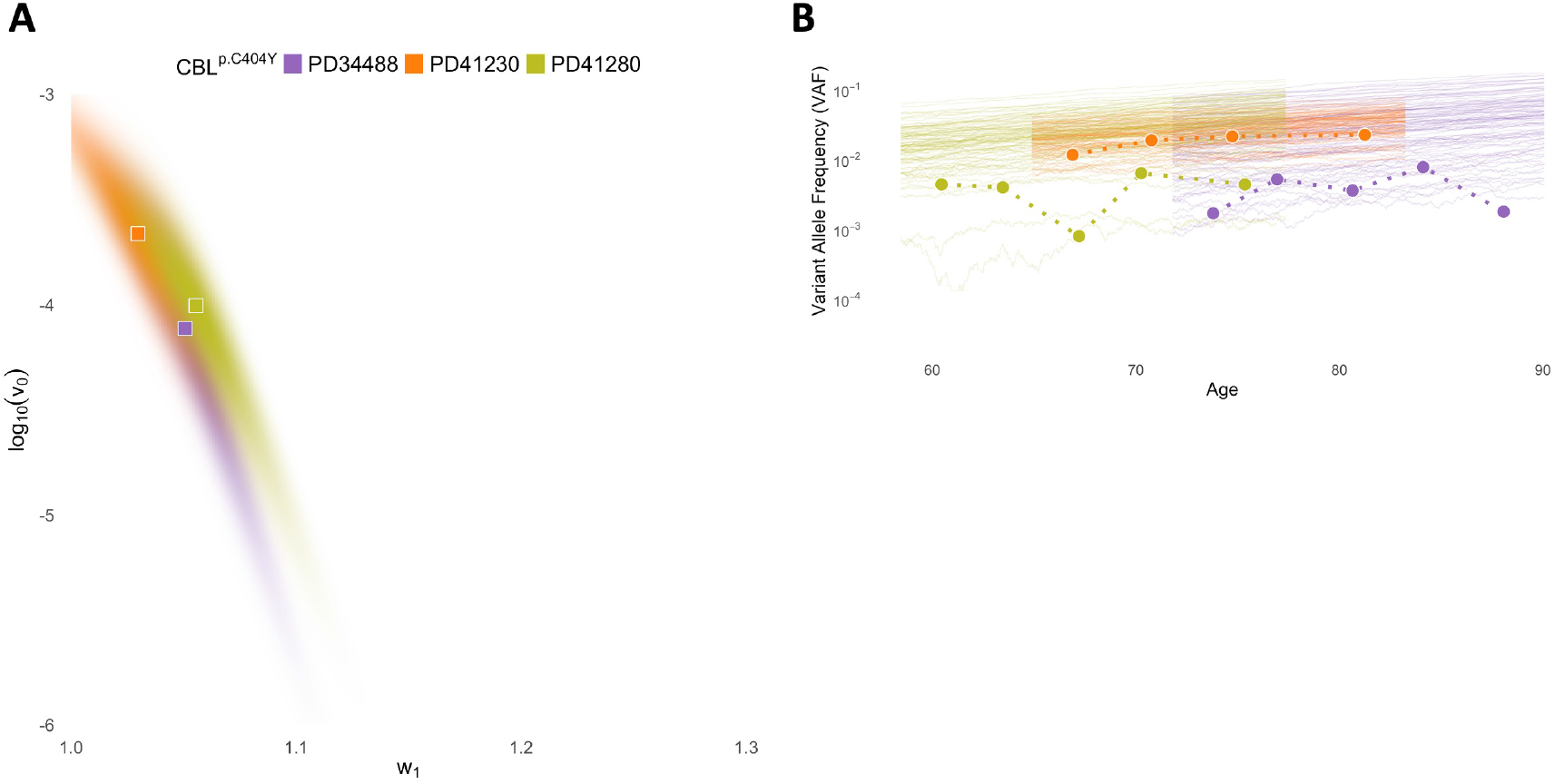
Inference of *CBL-C404Y* ‘s mutation rate and selection rate from time-series data in [Fabre et al., 2022]. **A**: Joint posterior distributions for log_10_(*v*_0_) and *w*_1_. Squares = MAP estimates. **B**: 100 simulated VAF trajectories in logscale assuming MAP estimates (thin lines), against observed VAFs (circles, connected by dashed lines). Colors in **B** correspond to datasets in **A**.

**Supplementary Figure 5:**
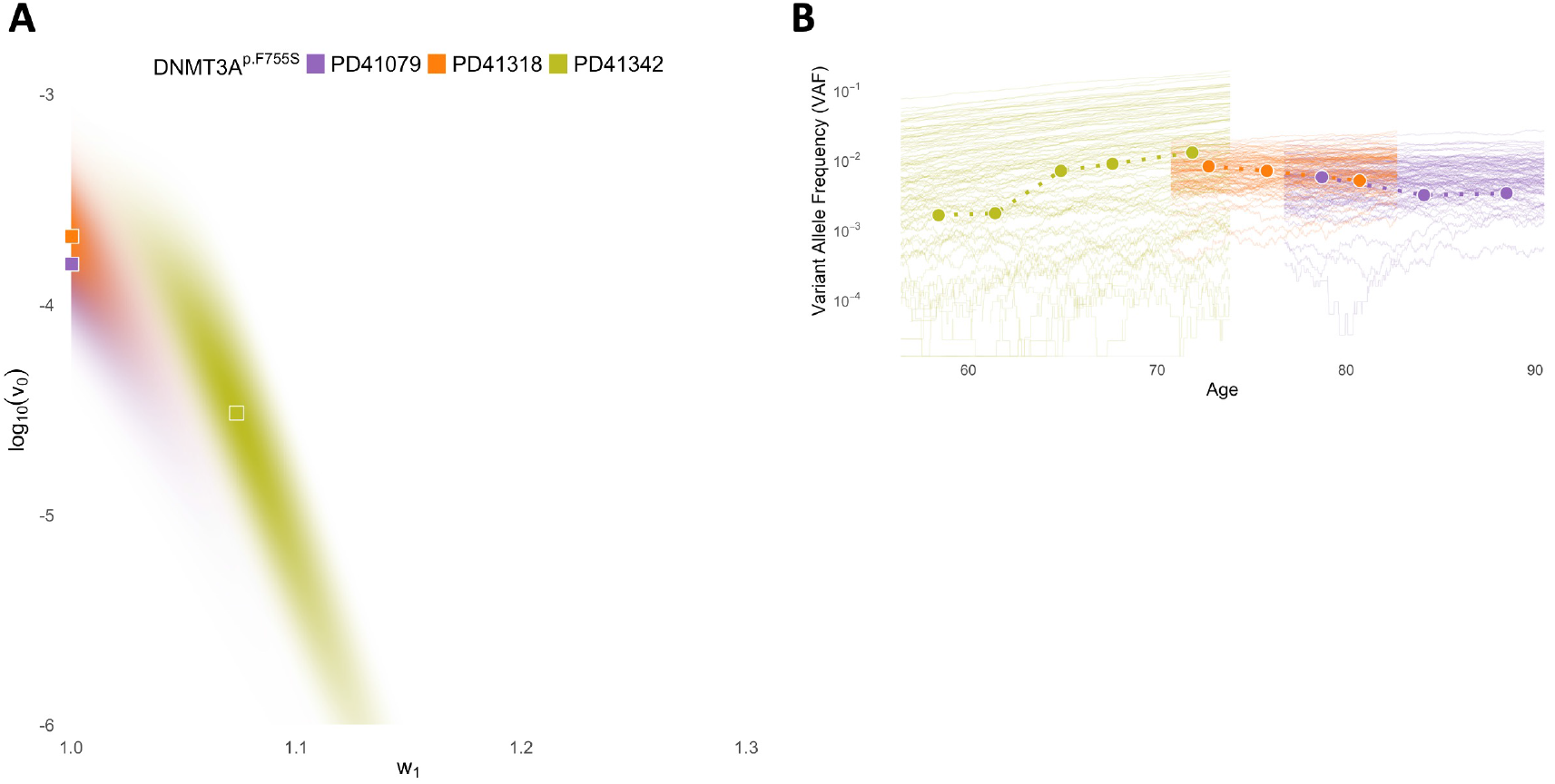
Inference of *DNMT3A-F755S* ‘s mutation rate and selection rate from time-series data in [Fabre et al., 2022]. **A**: Joint posterior distributions for log_10_(*v*_0_) and *w*_1_. Squares = MAP estimates. **B**: 100 simulated VAF trajectories in logscale assuming MAP estimates (thin lines), against observed VAFs (circles, connected by dashed lines). Colors in **B** correspond to datasets in **A**.

**Supplementary Figure 6:**
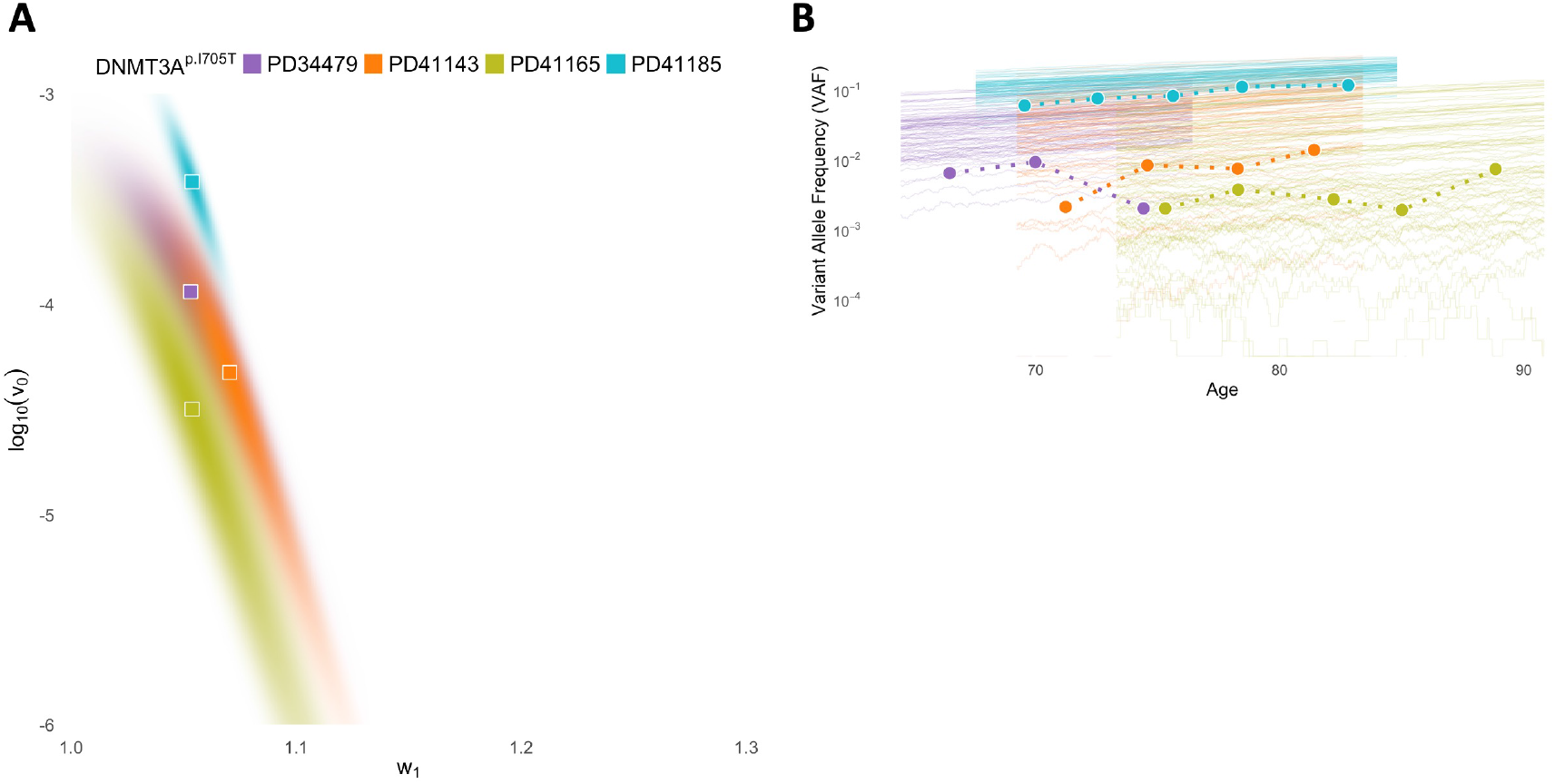
Inference of *DNMT3A-I705T* ‘s mutation rate and selection rate from time-series data in [Fabre et al., 2022]. **A**: Joint posterior distributions for log_10_(*v*_0_) and *w*_1_. Squares = MAP estimates. **B**: 100 simulated VAF trajectories in logscale assuming MAP estimates (thin lines), against observed VAFs (circles, connected by dashed lines). Colors in **B** correspond to datasets in **A**.

**Supplementary Figure 7:**
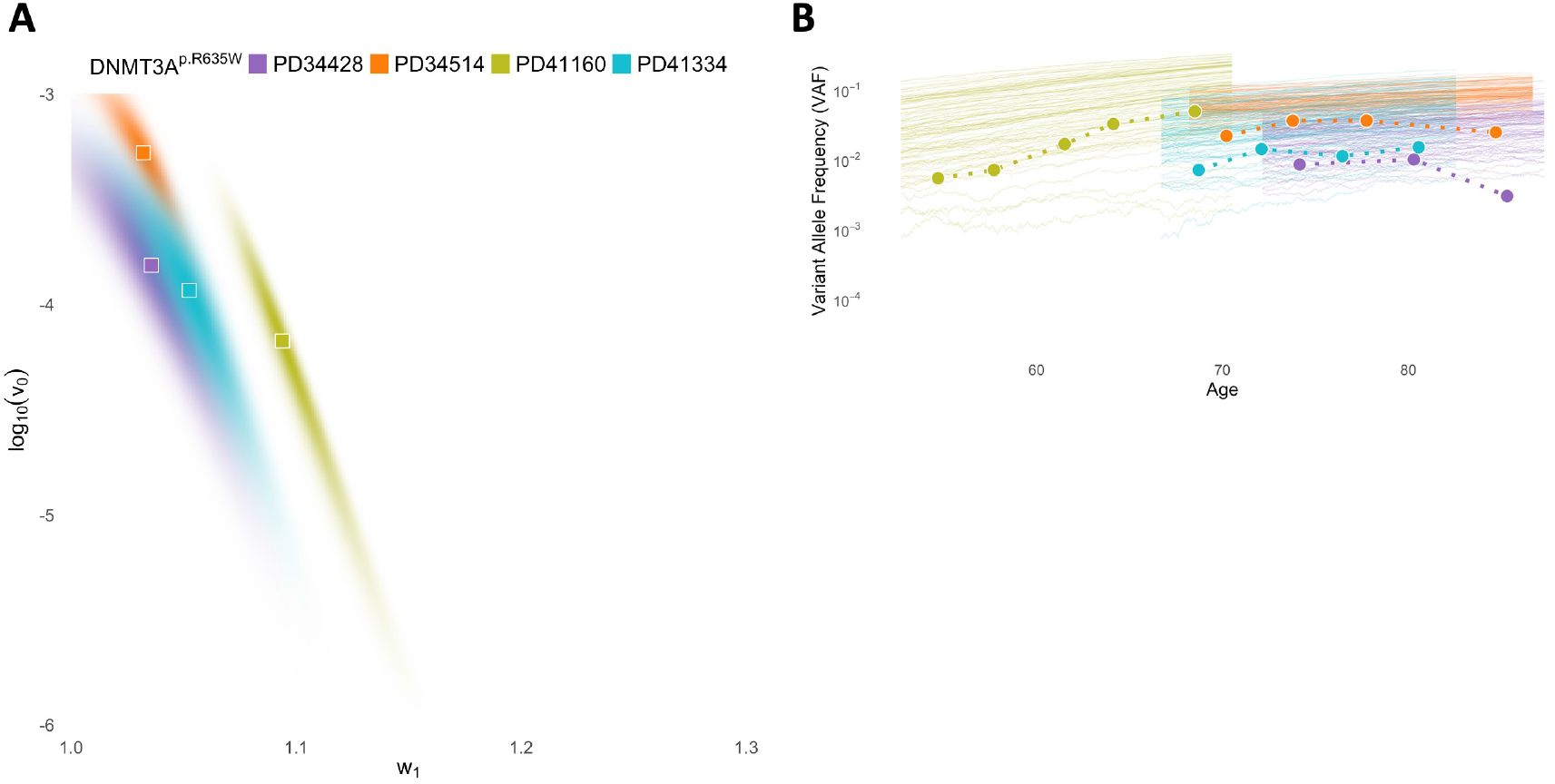
Inference of *DNMT3A-R635W* ‘s mutation rate and selection rate from time-series data in [Fabre et al., 2022]. **A**: Joint posterior distributions for log_10_(*v*_0_) and *w*_1_. Squares = MAP estimates. **B**: 100 simulated VAF trajectories in logscale assuming MAP estimates (thin lines), against observed VAFs (circles, connected by dashed lines). Colors in **B** correspond to datasets in **A**.

**Supplementary Figure 8:**
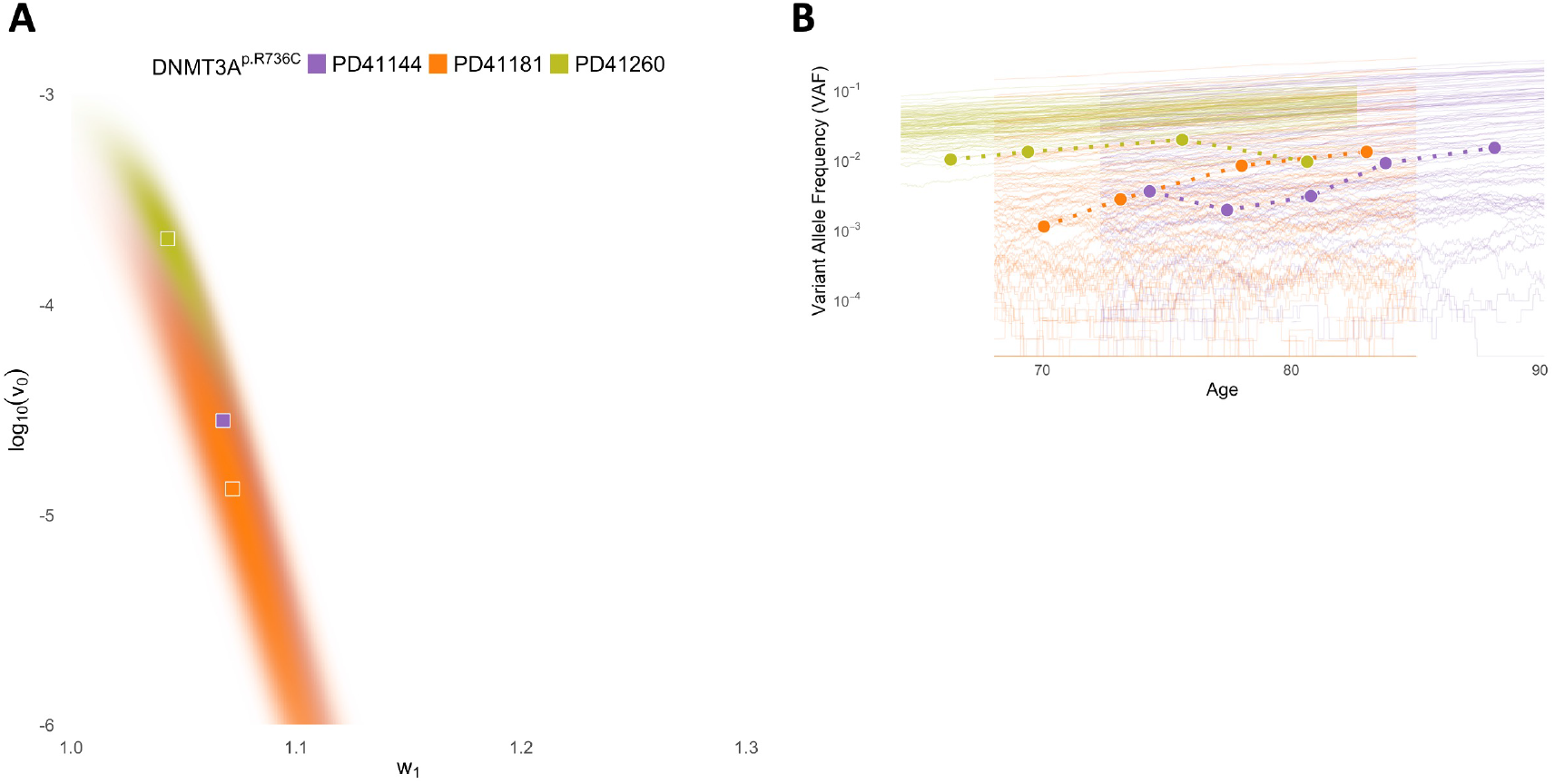
Inference of *DNMT3A-R736C* ‘s mutation rate and selection rate from time-series data in [Fabre et al., 2022]. **A**: Joint posterior distributions for log_10_(*v*_0_) and *w*_1_. Squares = MAP estimates. **B**: 100 simulated VAF trajectories in logscale assuming MAP estimates (thin lines), against observed VAFs (circles, connected by dashed lines). Colors in **B** correspond to datasets in **A**.

**Supplementary Figure 9:**
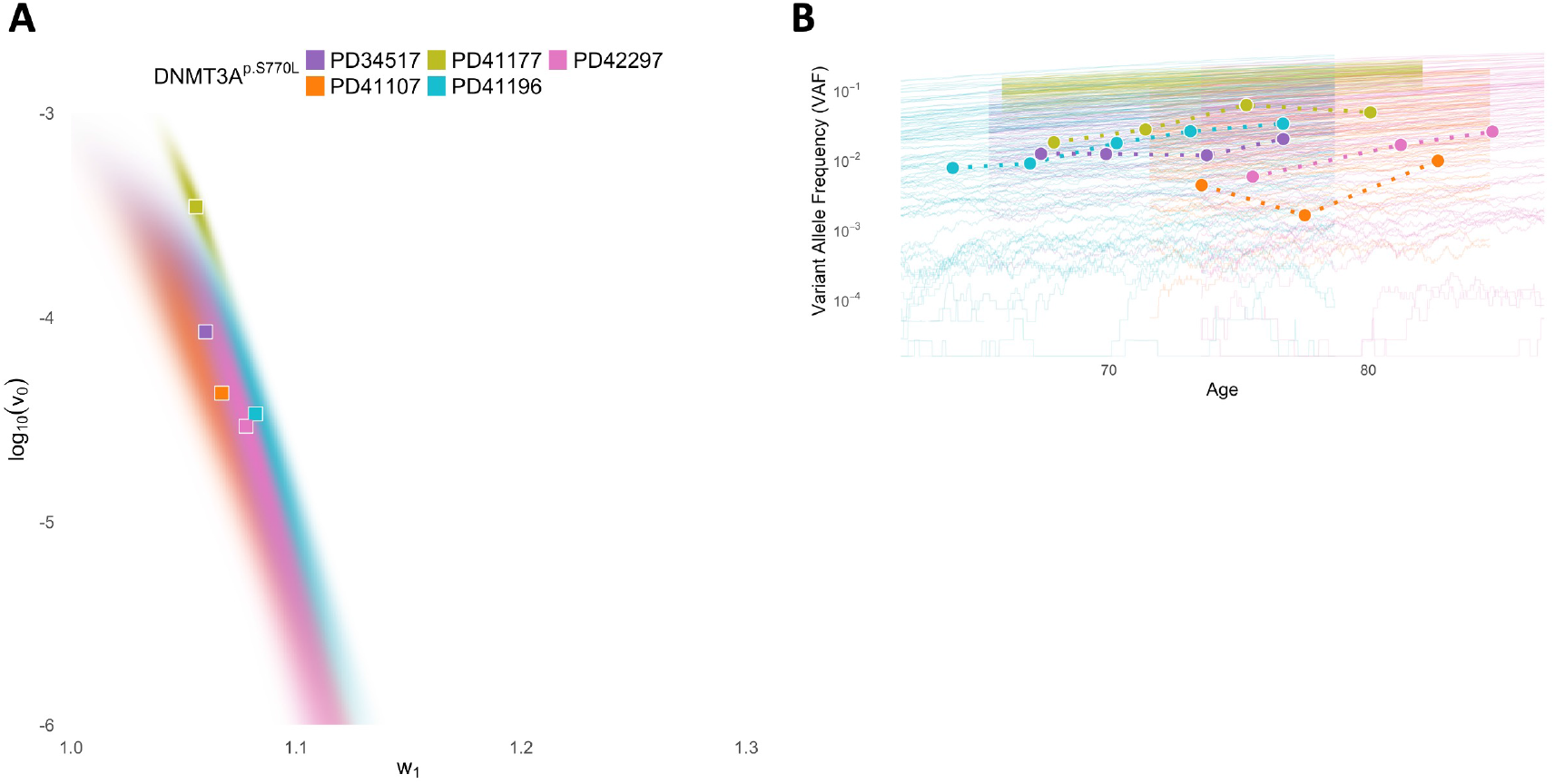
Inference of *DNMT3A-S770L*’s mutation rate and selection rate from time-series data in [Fabre et al., 2022]. **A**: Joint posterior distributions for log_10_(*v*_0_) and *w*_1_. Squares = MAP estimates. **B**: 100 simulated VAF trajectories in logscale assuming MAP estimates (thin lines), against observed VAFs (circles, connected by dashed lines). Colors in **B** correspond to datasets in **A**.

**Supplementary Figure 10:**
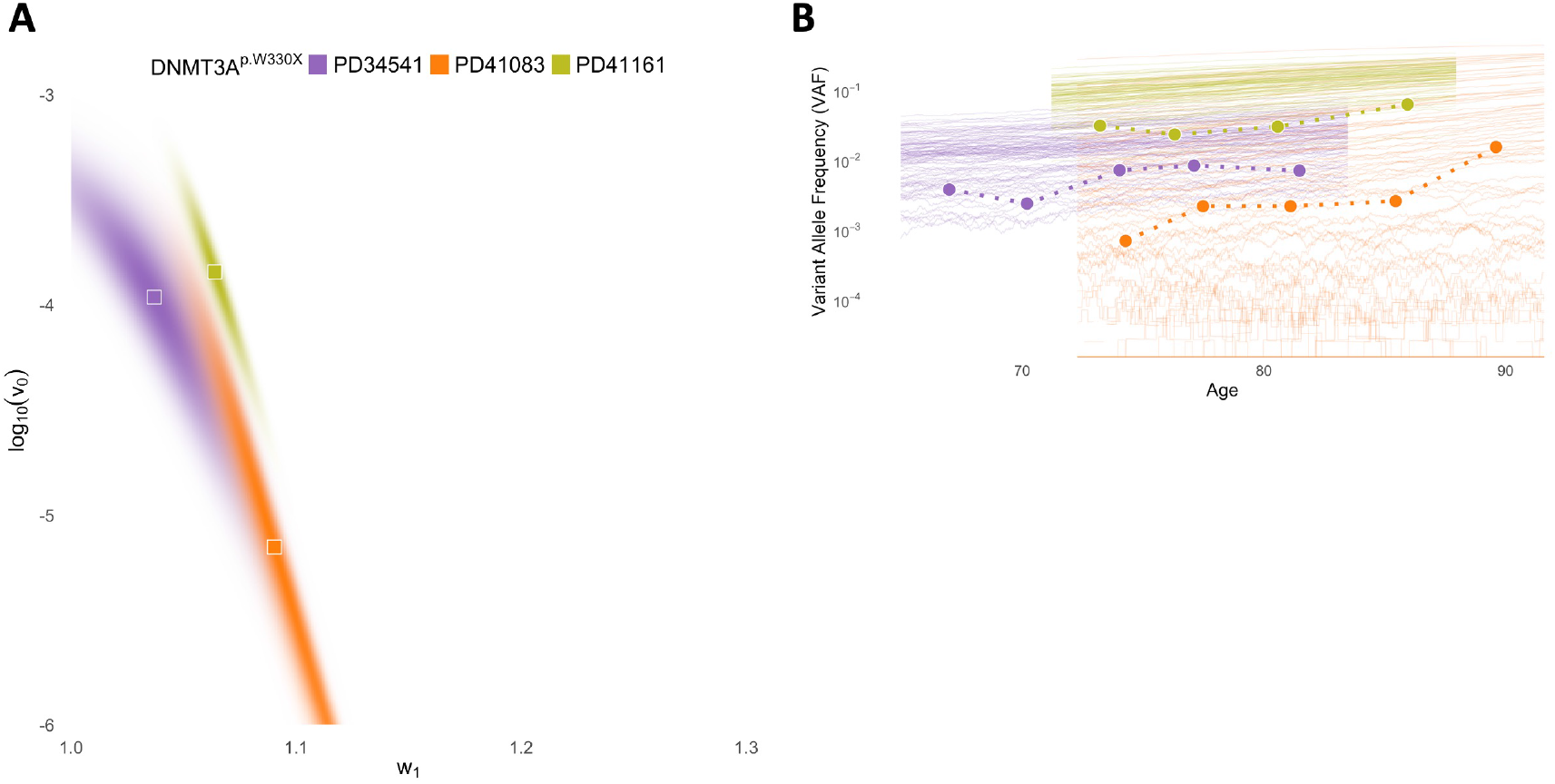
Inference of *DNMT3A-W330X* ‘s mutation rate and selection rate from time-series data in [Fabre et al., 2022]. **A**: Joint posterior distributions for log_10_(*v*_0_) and *w*_1_. Squares = MAP estimates. **B**: 100 simulated VAF trajectories in logscale assuming MAP estimates (thin lines), against observed VAFs (circles, connected by dashed lines). Colors in **B** correspond to datasets in **A**.

**Supplementary Figure 11:**
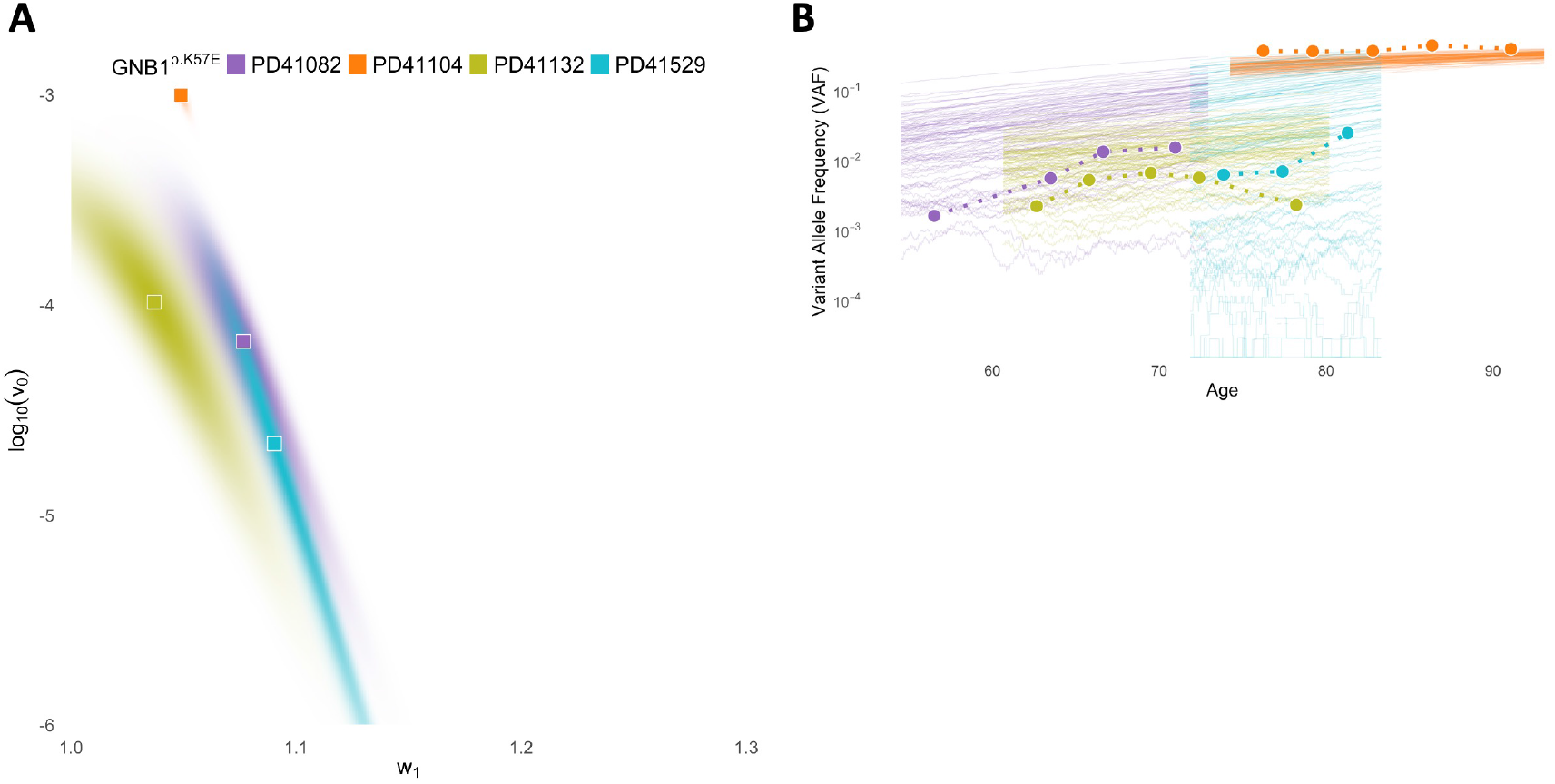
Inference of *GNB1-K57E* ‘s mutation rate and selection rate from time-series data in [Fabre et al., 2022]. **A**: Joint posterior distributions for log_10_(*v*_0_) and *w*_1_. Squares = MAP estimates. **B**: 100 simulated VAF trajectories in logscale assuming MAP estimates (thin lines), against observed VAFs (circles, connected by dashed lines). Colors in **B** correspond to datasets in **A**.

**Supplementary Figure 12:**
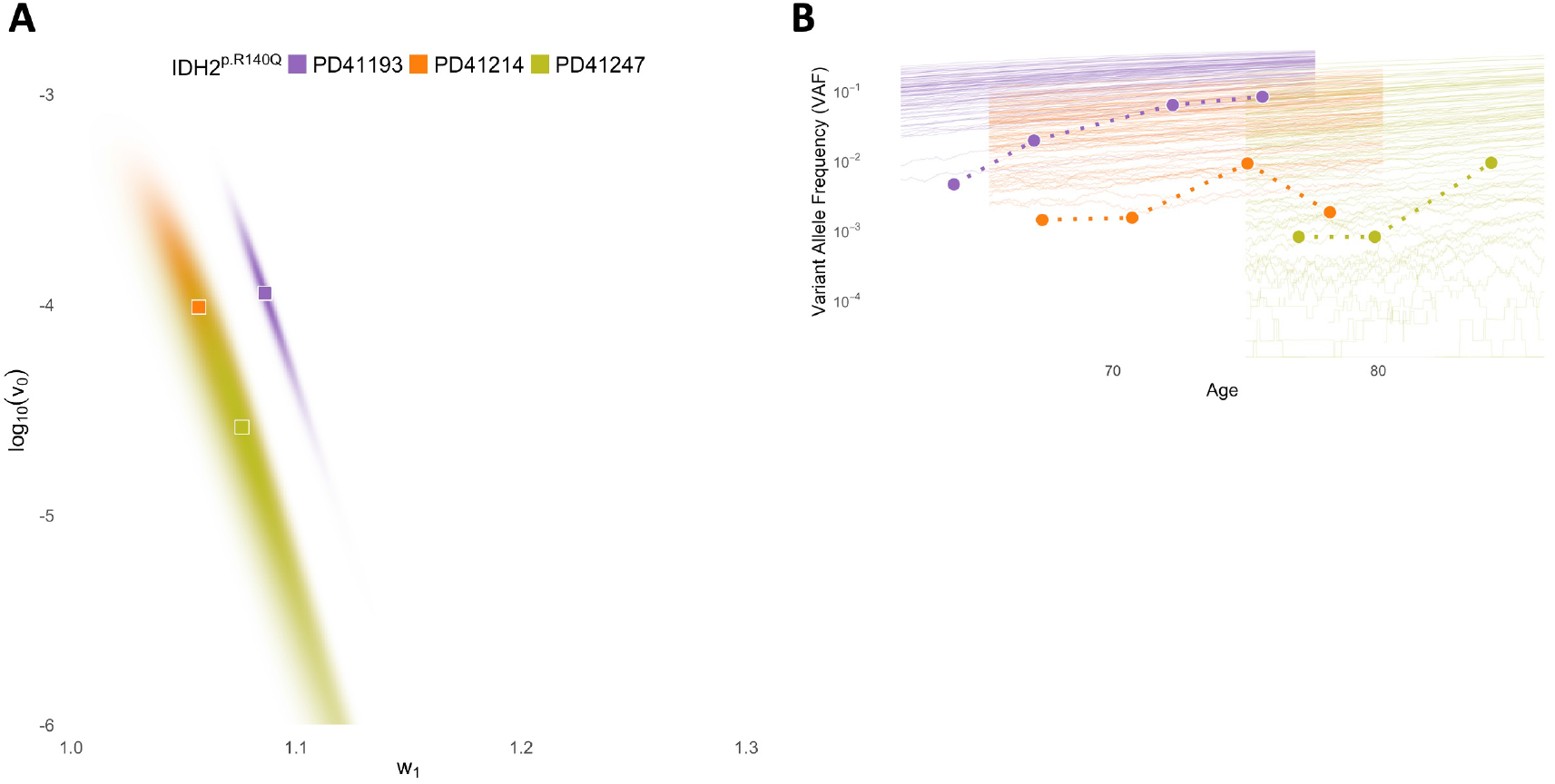
Inference of *IDH2-R140Q* ‘s mutation rate and selection rate from time-series data in [Fabre et al., 2022]. **A**: Joint posterior distributions for log_10_(*v*_0_) and *w*_1_. Squares = MAP estimates. **B**: 100 simulated VAF trajectories in logscale assuming MAP estimates (thin lines), against observed VAFs (circles, connected by dashed lines). Colors in **B** correspond to datasets in **A**.

**Supplementary Figure 13:**
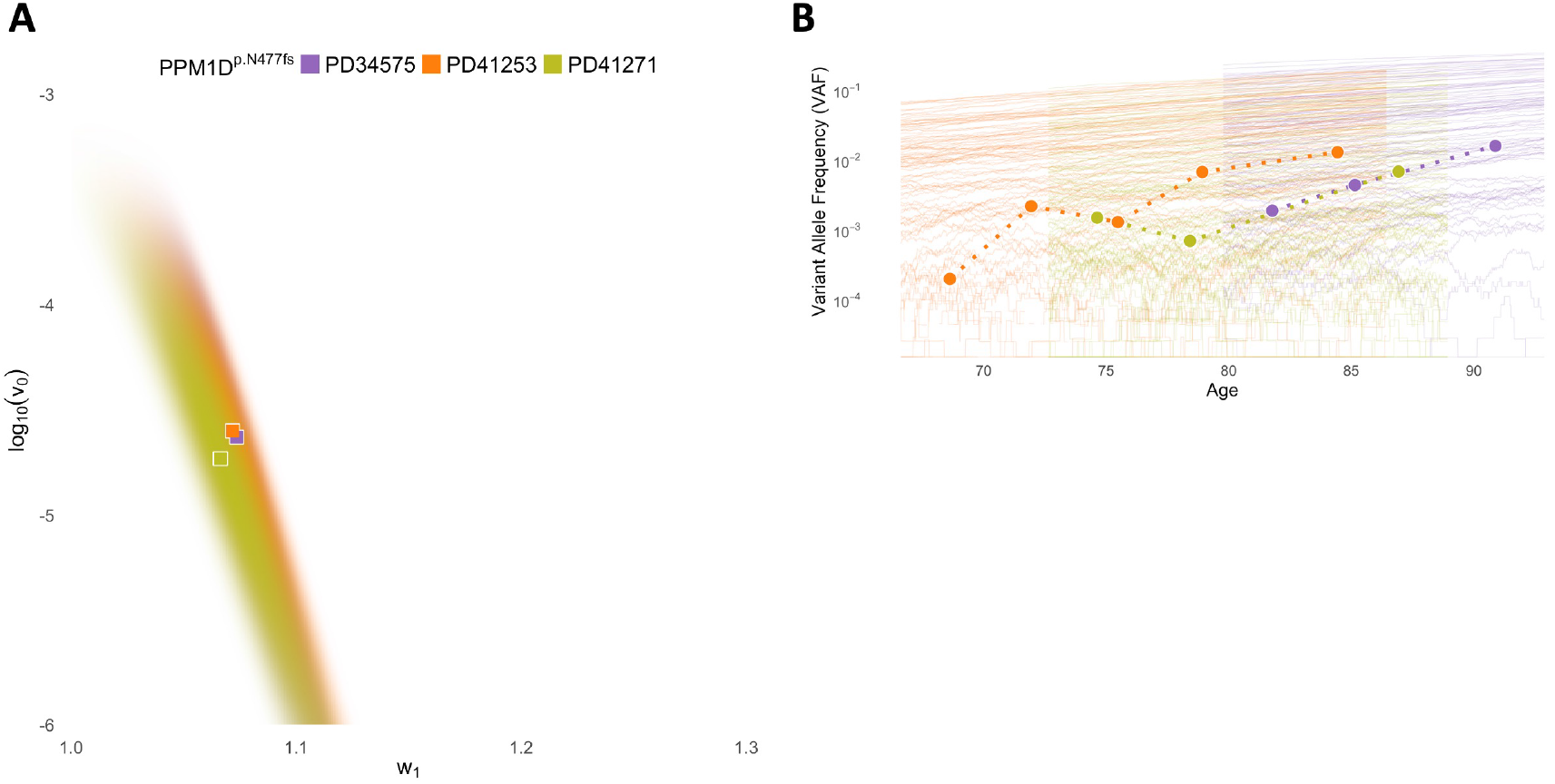
Inference of *PPM1D-N477fs*’s mutation rate and selection rate from time-series data in [Fabre et al., 2022]. **A**: Joint posterior distributions for log_10_(*v*_0_) and *w*_1_. Squares = MAP estimates. **B**: 100 simulated VAF trajectories in logscale assuming MAP estimates (thin lines), against observed VAFs (circles, connected by dashed lines). Colors in **B** correspond to datasets in **A**.

**Supplementary Figure 14:**
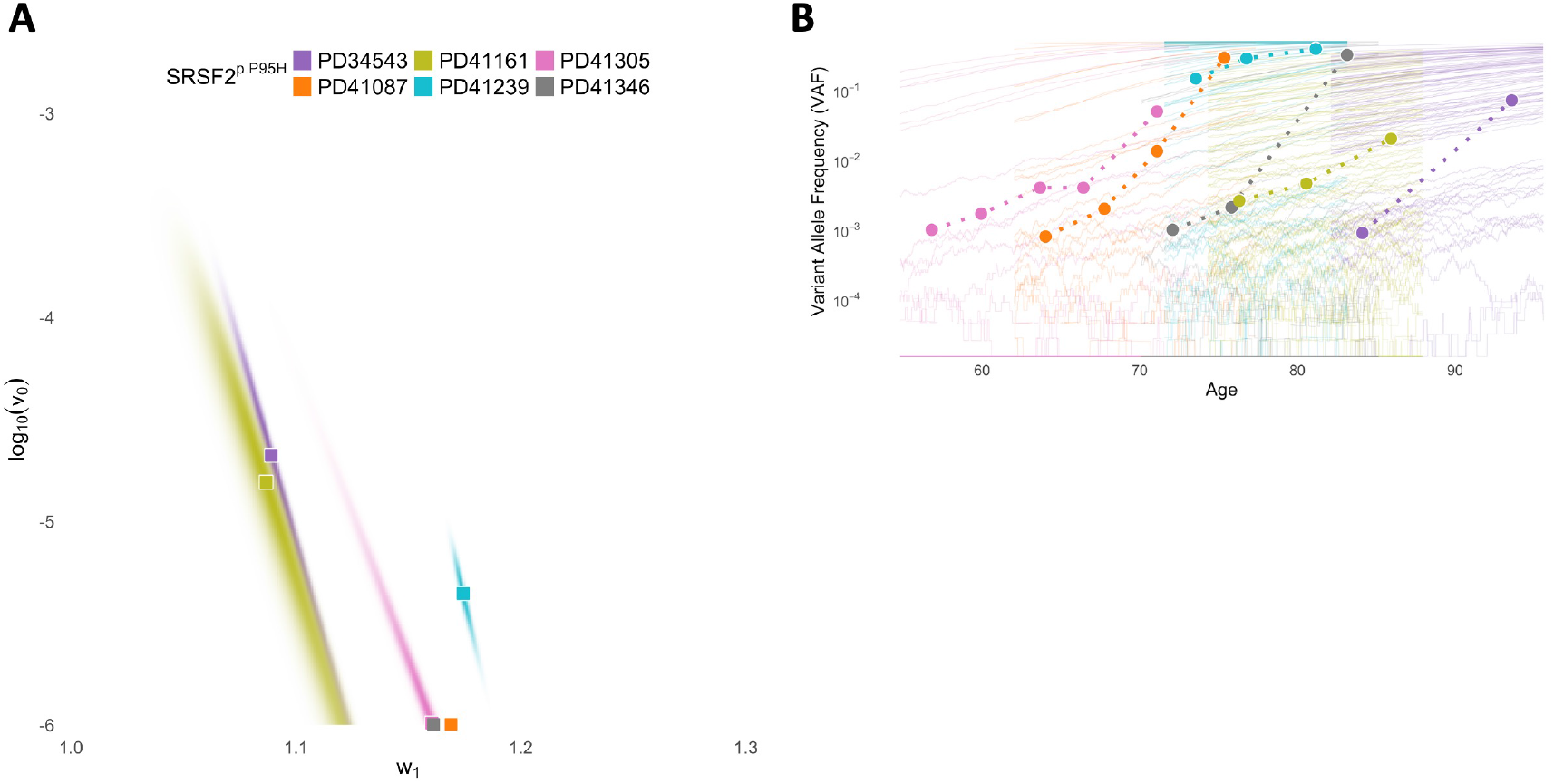
Inference of *SRSF2-P95H* ‘s mutation rate and selection rate from time-series data in [Fabre et al., 2022]. **A**: Joint posterior distributions for log_10_(*v*_0_) and *w*_1_. Squares = MAP estimates. **B**: 100 simulated VAF trajectories in logscale assuming MAP estimates (thin lines), against observed VAFs (circles, connected by dashed lines). Colors in **B** correspond to datasets in **A**.

**Supplementary Figure 15:**
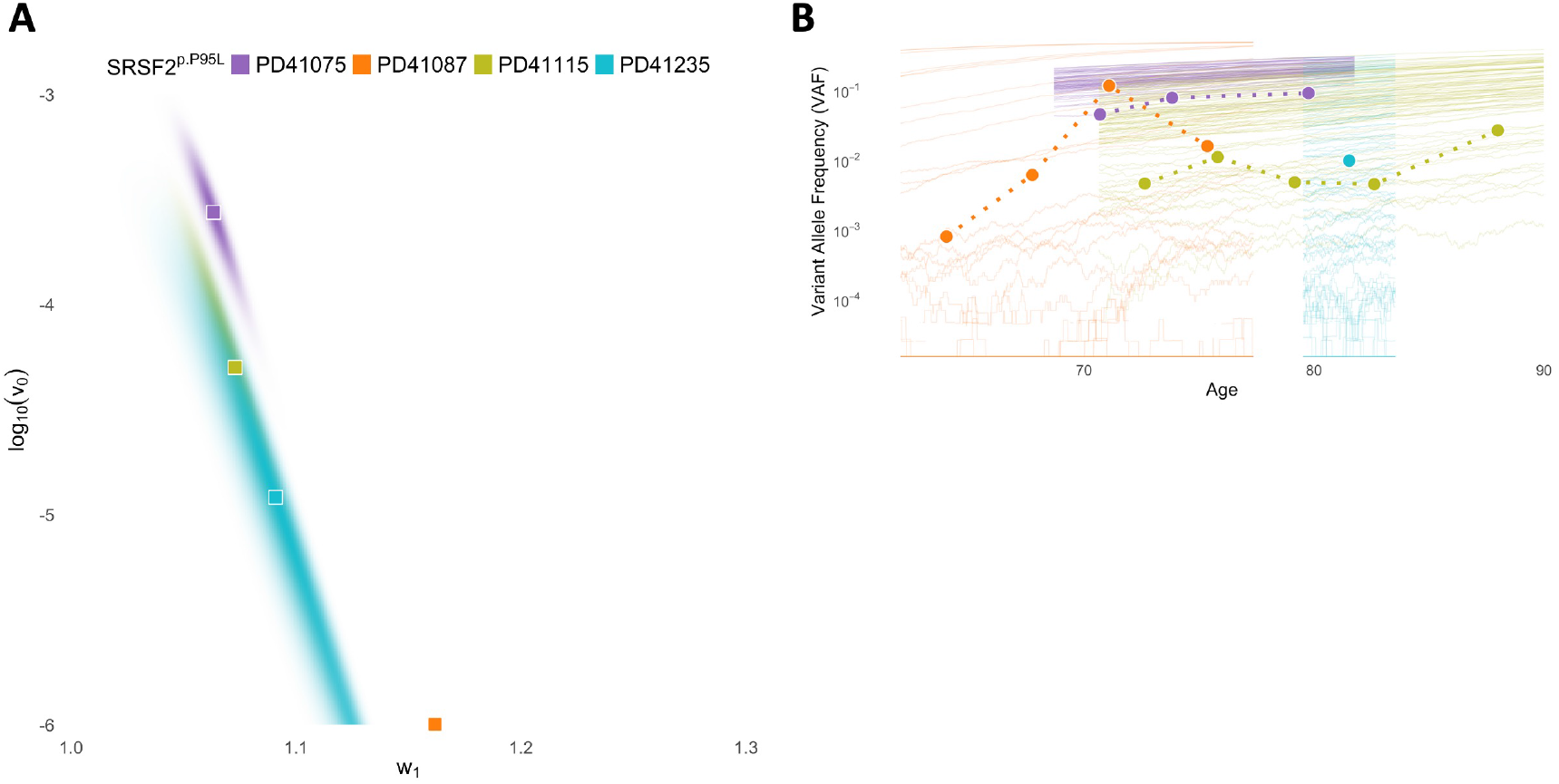
Inference of *SRSF2-P95L*’s mutation rate and selection rate from time-series data in [Fabre et al., 2022]. **A**: Joint posterior distributions for log_10_(*v*_0_) and *w*_1_. Squares = MAP estimates. **B**: 100 simulated VAF trajectories in logscale assuming MAP estimates (thin lines), against observed VAFs (circles, connected by dashed lines). Colors in **B** correspond to datasets in **A**.

**Supplementary Figure 16:**
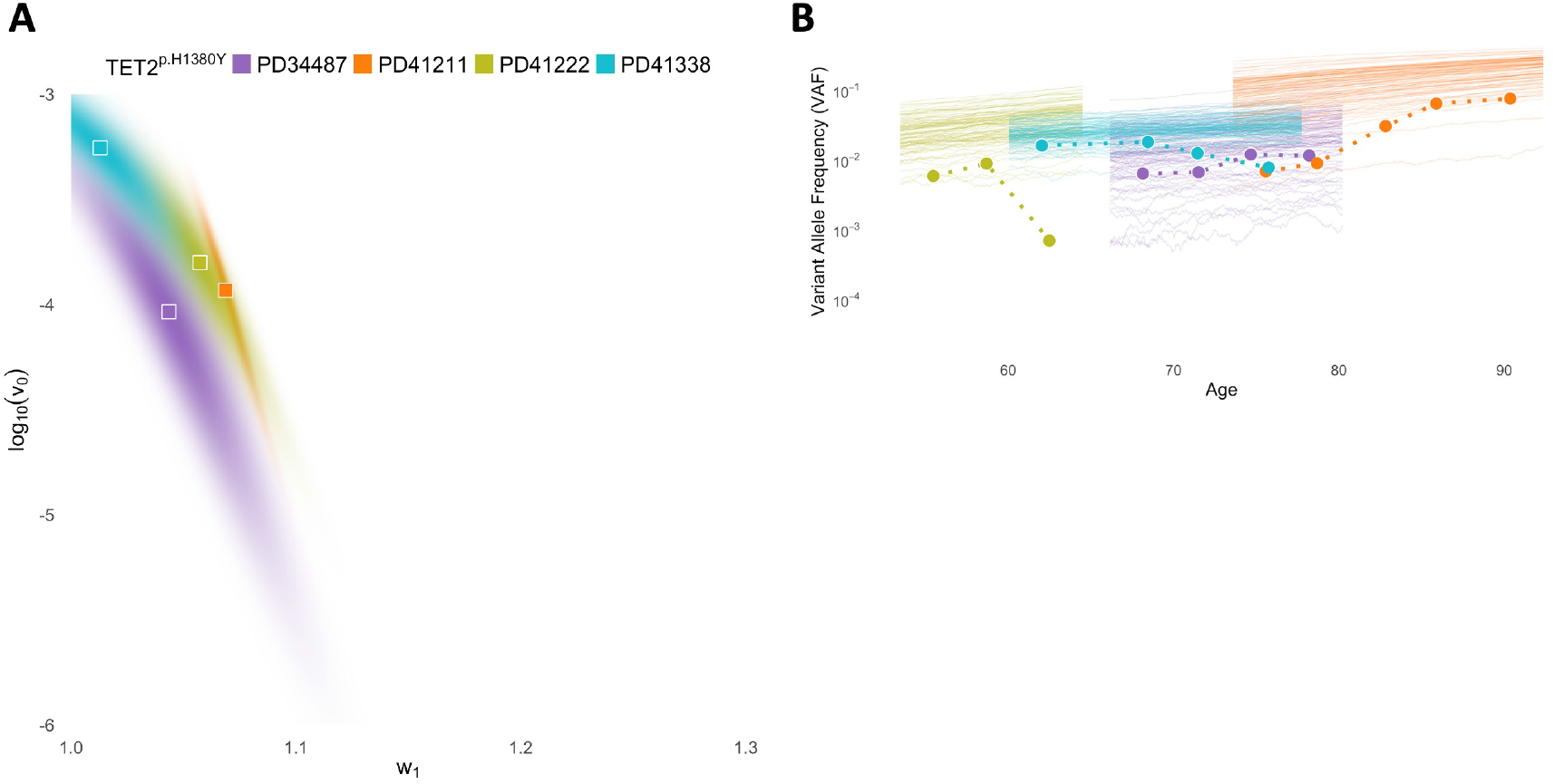
Inference of *TET2-H1380Y* ‘s mutation rate and selection rate from time-series data in [Fabre et al., 2022]. **A**: Joint posterior distributions for log_10_(*v*_0_) and *w*_1_. Squares = MAP estimates. **B**: 100 simulated VAF trajectories in logscale assuming MAP estimates (thin lines), against observed VAFs (circles, connected by dashed lines). Colors in **B** correspond to datasets in **A**.

**Supplementary Figure 17:**
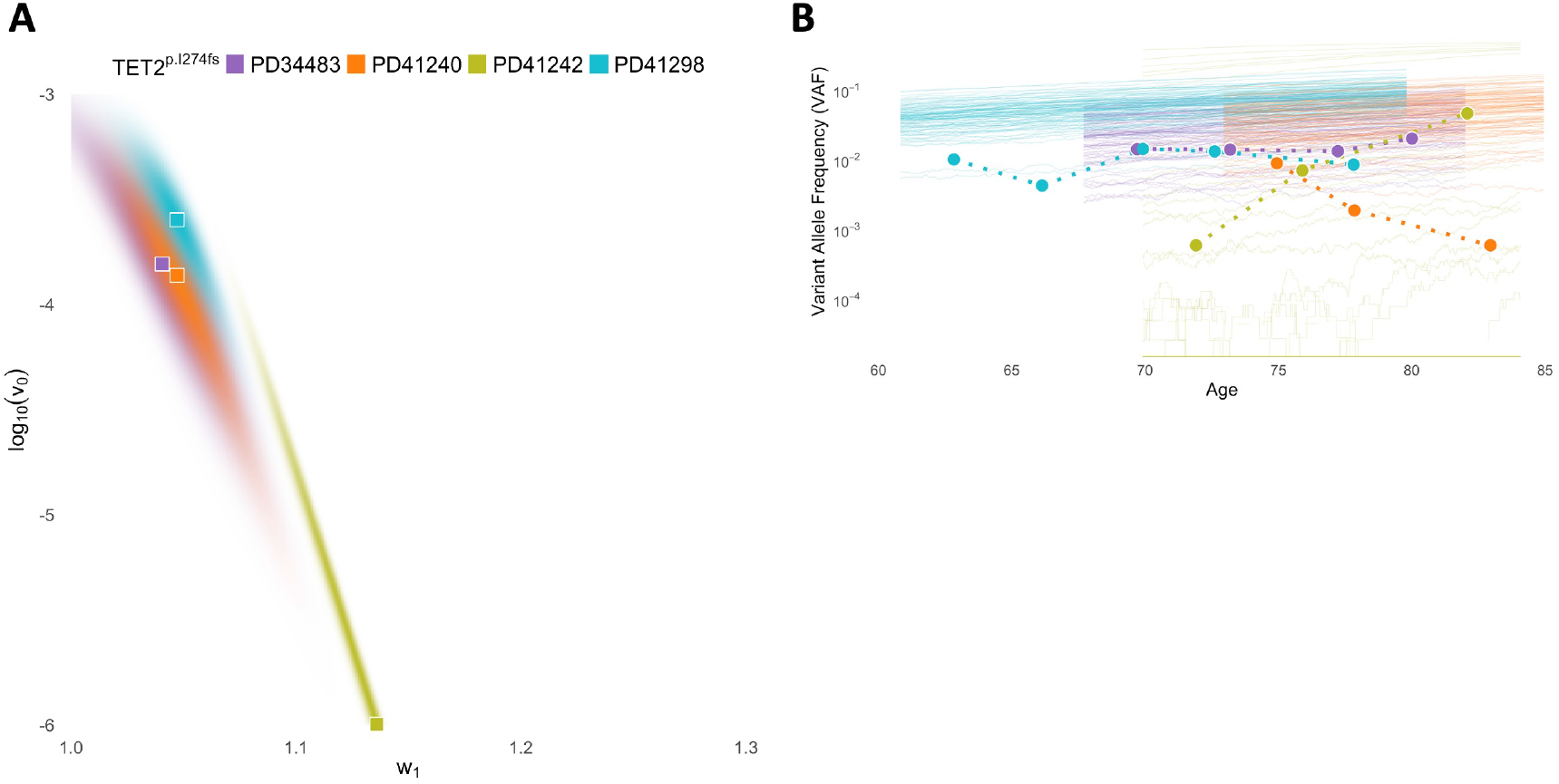
Inference of *TET2-I274fs*’s mutation rate and selection rate from time-series data in [Fabre et al., 2022]. **A**: Joint posterior distributions for log_10_(*v*_0_) and *w*_1_. Squares = MAP estimates. **B**: 100 simulated VAF trajectories in logscale assuming MAP estimates (thin lines), against observed VAFs (circles, connected by dashed lines). Colors in **B** correspond to datasets in **A**.

**Supplementary Figure 18:**
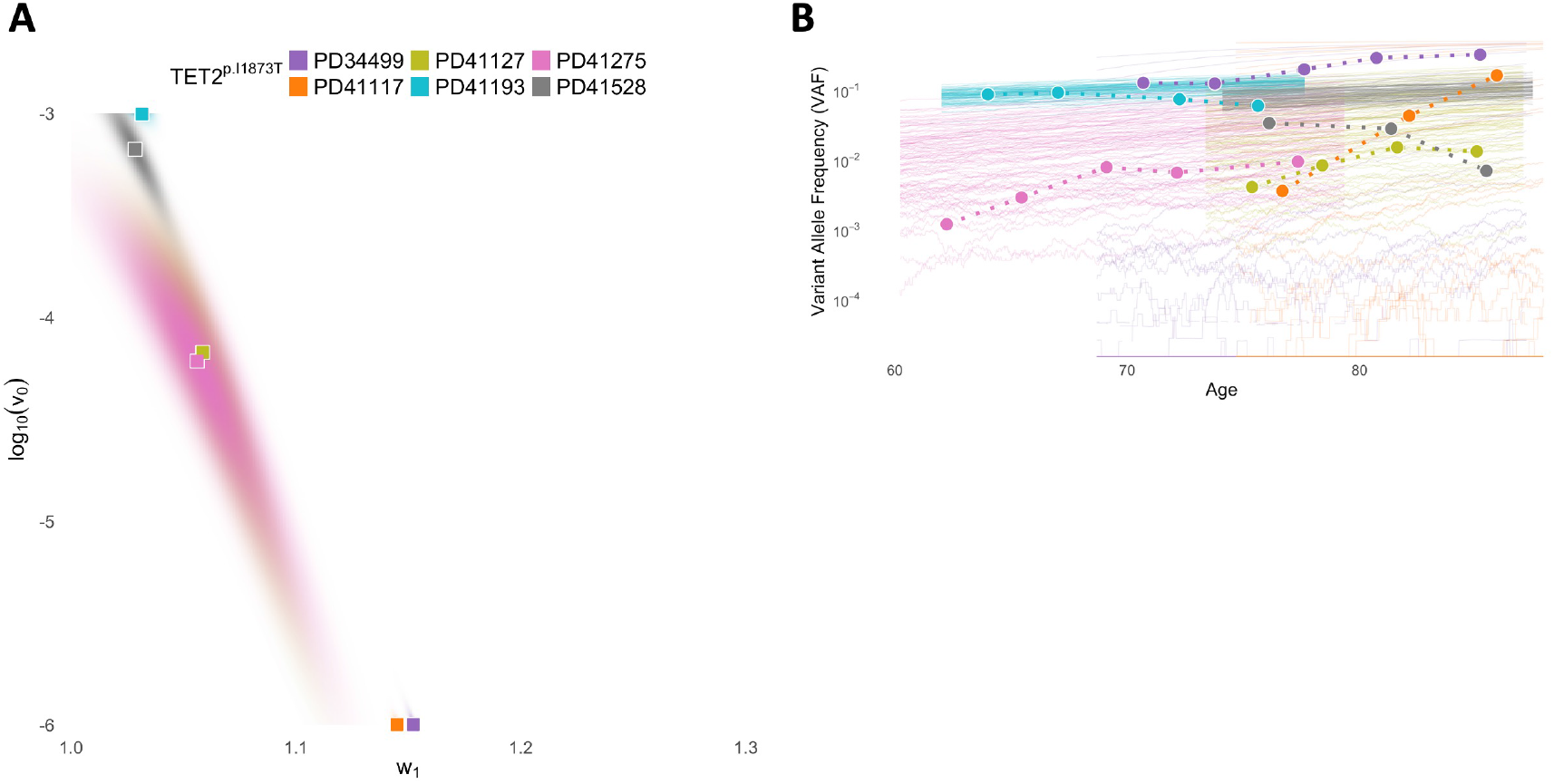
Inference of *TET2-I1873T* ‘s mutation rate and selection rate from time-series data in [Fabre et al., 2022]. **A**: Joint posterior distributions for log_10_(*v*_0_) and *w*_1_. Squares = MAP estimates. **B**: 100 simulated VAF trajectories in logscale assuming MAP estimates (thin lines), against observed VAFs (circles, connected by dashed lines). Colors in **B** correspond to datasets in **A**.

**Supplementary Figure 19:**
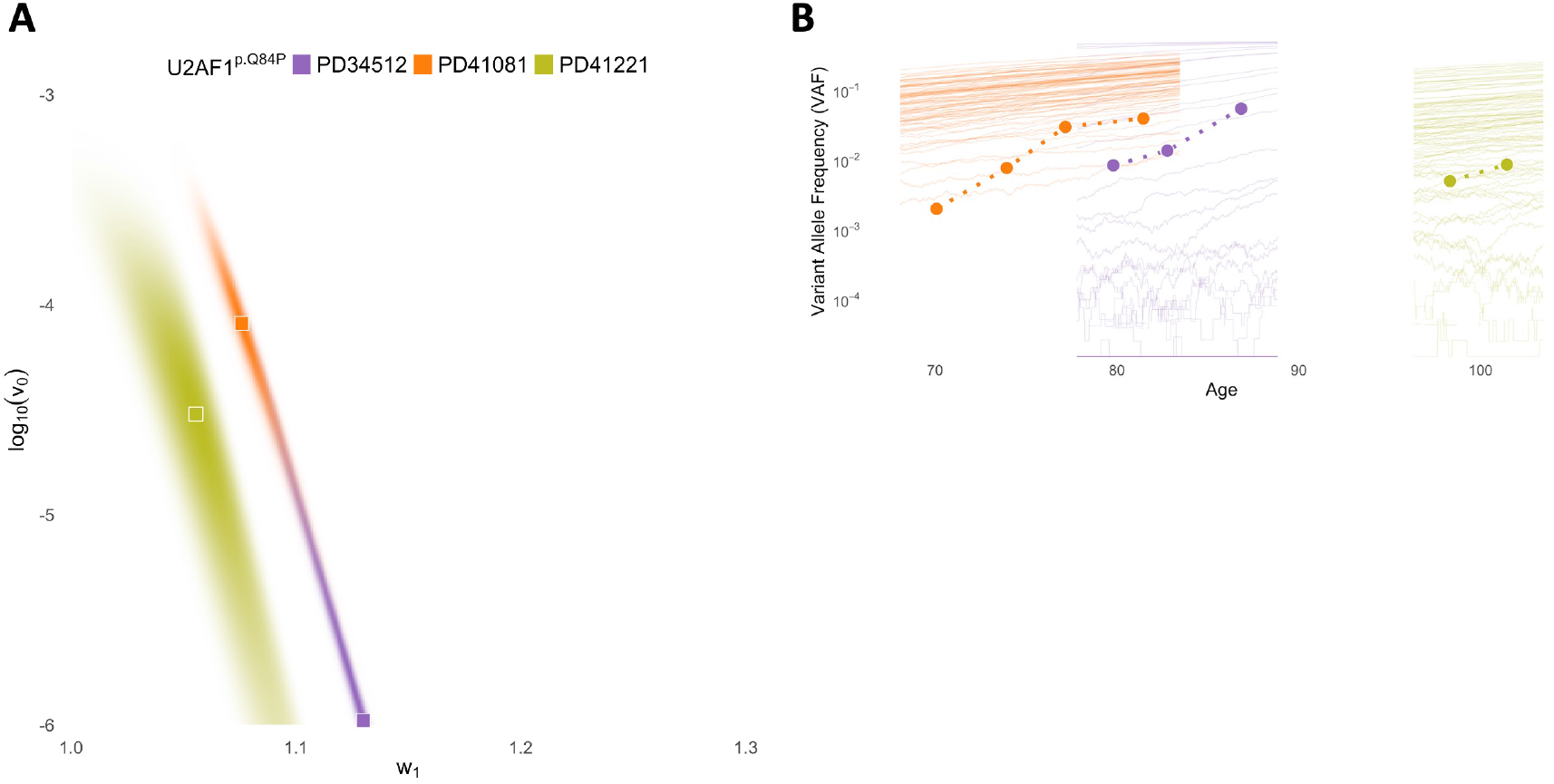
Inference of *U2AF1-Q84P* ‘s mutation rate and selection rate from time-series data in [Fabre et al., 2022]. **A**: Joint posterior distributions for log_10_(*v*_0_) and *w*_1_. Squares = MAP estimates. **B**: 100 simulated VAF trajectories in logscale assuming MAP estimates (thin lines), against observed VAFs (circles, connected by dashed lines). Colors in **B** correspond to datasets in **A**.

## A Appendix

### A.1 Construction of Dynamics

Let 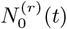 and 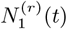 denote the number of WT (type 0) and CH (type 1) cells at time *t* with initial population consisting of *r* WT cells. We denote events in the dynamics by (*j, a*), where *j* ∈ {0, 1} and *a* ∈ {*b, m, d*} (birth, mutation, and death). Event (*j, a*) refers to a type *j* individual taking action *a*. We construct the model following the Poisson process formulation in [Ethier and Kurtz, 2009]. Let 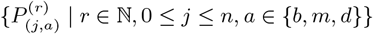 be a set of independent rate 1 Poisson processes.

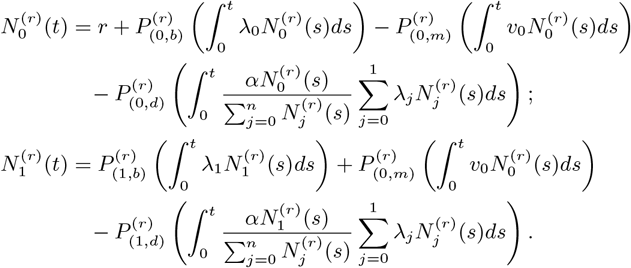

For all 0 ≤ *j* ≤ 1, division rate *λ*_*j*_ *>* 0 and mutation rate *v*_0_ *>* 0. Since type 1 individuals cannot further mutate, *v*_1_ = 0. For our model, we define fitness of a type *j* individual by *w*_*j*_ := *λ*_*j*_ − *v*_*j*_ .

### B FLLN and FCLT

#### Proposition 1.

*As r* → ∞, 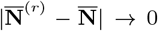 *almost surely in Skorokhod space* (𝔻 ([0, ∞)), *d*_∞_), *where* 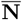 *is characterized by the following autonomous system of ODEs:*

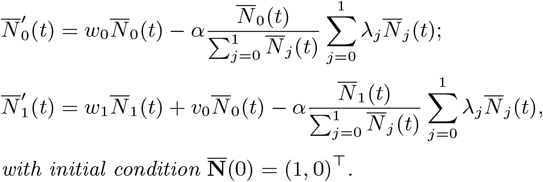

*Proof*. We verify conditions of Theorem 2.1 in Ethier and Kurtz [2009]. Rewrite the system of differential equations as

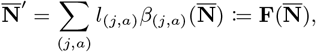

where *l*_(*j,a*)_ is a vector representing the change of population and *β*_(*j,a*)_ is the rate for that event. For events associated with type 0 indiduals,

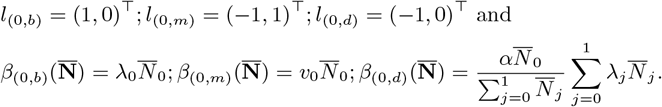

For events associated with type 1 individuals,

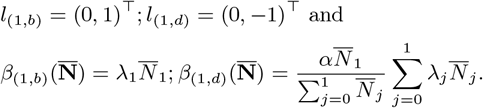

Since we only have finitely many events and *β*_(*j,a*)_’s are continuous, we have for every compact *K* ⊂ (0, ∞)^2^,

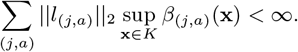

Since *β*_(*j,b*)_, *β*_(*j,m*)_, and *β*_(*j,d*)_ are continuously differentiable on (0, ∞)^2^, **F** is locally Lipschitz. Hence, by Theorem 2.1 in Ethier and Kurtz [2009], we have as *r* → ∞,

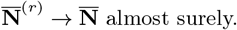

In the proof, we omit one state (0, 0)^⊤^ as it has probability tending to zero as *r* → ∞.

#### Proposition 2.

*As r* → ∞, 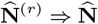, *where* 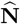 *satisfies*

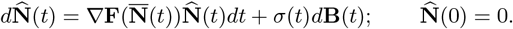

**B** *is a vector of standard Brownian motions and σ*(*t*) *is a* 2 *by* 5 *matrix. The expression for σ*(*t*) *is*

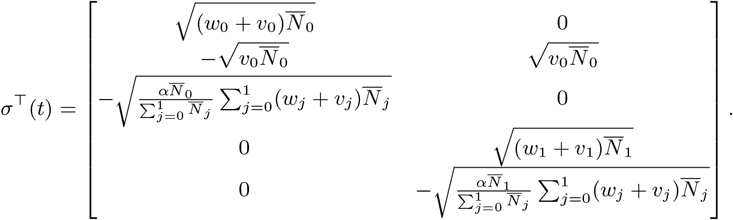

*Proof*. We verify conditions in Theorem 2.3 on page 458 of Ethier and Kurtz [2009]. Recall *l*_(*j,a*)_’s and *β*_(*j,a*)_’s defined in Proposition 1. By continuity of *β*_(*j,a*)_’s, we have for every compact *K* ⊂ (0, ∞)^2^,

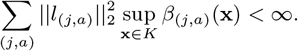

Since *β*_(*j,a*)_ is continuously differentiable for all (*j, a*)’s, ∇**F** is continuous. Therefore, 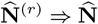 as *r* → ∞, where 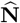 is defined by the stochastic differential equation in the statement of this Proposition.

### C Ratio Dynamics

For *t* ≥ 0, we define the ratio dynamics as

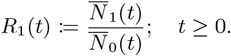

#### Lemma 1.

*For t* ≥ 0, *we have*

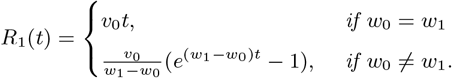

*Proof*. From 1, we have

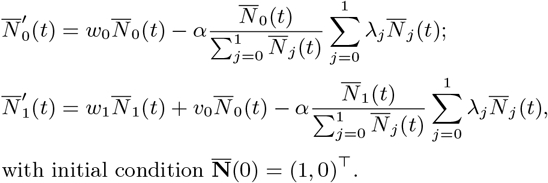

Therefore,

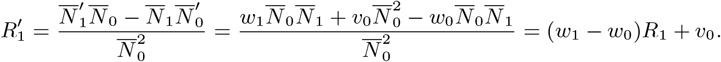

This completes the proof

### D Approximated Dynamics of VAF

For *t* ≥ 0, we define the VAF dynamics as

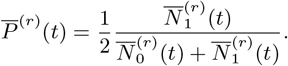

#### Theorem 1.

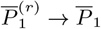 *almost surely, where*

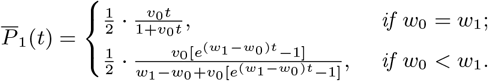

*Proof*. We define *g* : 𝒟^2^ → 𝒟 such that for all *x, y* ∈ 𝒟,

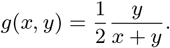

It is evident that *g*(*x, y*) is an element of 𝒟. Usually, the addition operator is not continuous on space 𝒟. That is, there exists *x*_*n*_ → *x* and *y*_*n*_ → *y* in *D*, but *x*_*n*_ + *y*_*n*_ ↛ *x* + *y*. For a concrete example, see Example 3.3.1 in Whitt [2002]. Nonetheless, if both *x* and *y* are continuous, we do have lim_*n*→∞_(*x*_*n*_ + *y*_*n*_) = *x* + *y*. Using similar reasoning, if both *x* and *y* are continuous, the ratio operator is also continuous. Hence, *g*(*x*_*n*_, *y*_*n*_) converges to *g*(*x, y*) in ∥ · ∥_*T*_ for all *T >* 0. This concludes that *g* is continuous at 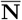 and by the continuous mapping theorem, we have

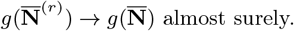

#### Theorem 2.

*As r* → ∞, *the finite dimensional distributions of* 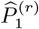 *converges weakly to the finite dimensional distribution of* 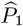 *defined by*

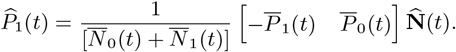

*Proof*. This is demonstrated by the delta method. Since as *r* → ∞,

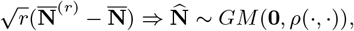

we can derive its finite dimensional distribution on (*t*_1_, · · ·, *t*_*n*_), which is a multivariate Gaussian distribution with mean being the zero vector and the covariance matrix *S* being

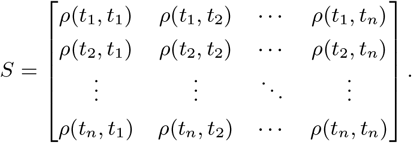

Define a function *h* : ℝ^2*n*^ → ℝ^*n*^ such that

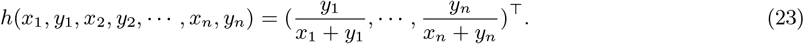

Therefore, the Jacobian of *h* is

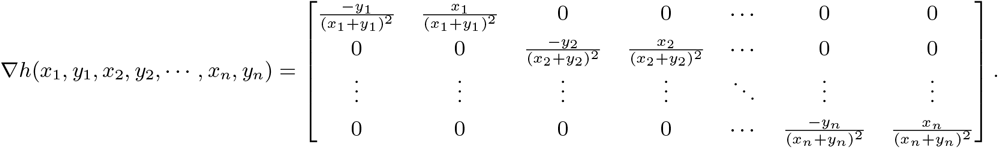

By delta method, we have

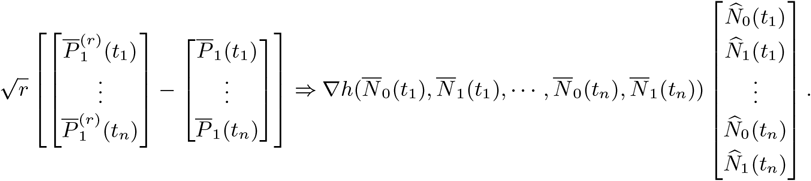

Hence, finite dimensional distribution of 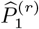 converges to that 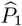.

